# Secondary metabolism drives ecological breadth in the Xylariaceae

**DOI:** 10.1101/2021.06.01.446356

**Authors:** Mario E.E. Franco, Jennifer H. Wisecaver, A. Elizabeth Arnold, Yu-Ming Ju, Jason C. Slot, Steven Ahrendt, Lillian P. Moore, Katharine E. Eastman, Kelsey Scott, Zachary Konkel, Stephen J. Mondo, Alan Kuo, Richard Hayes, Sajeet Haridas, Bill Andreopoulos, Robert Riley, Kurt LaButti, Jasmyn Pangilinan, Anna Lipzen, Mojgan Amirebrahimi, Juying Yan, Catherine Adam, Keykhosrow Keymanesh, Vivian Ng, Katherine Louie, Trent Northen, Elodie Drula, Bernard Henrissat, Huei-Mei Hsieh, Ken Youens-Clark, François Lutzoni, Jolanta Miadlikowska, Daniel C. Eastwood, Richard C. Hamelin, Igor V. Grigoriev, Jana M. U’Ren

## Abstract

Global, large-scale surveys of phylogenetically diverse plant and lichen hosts have revealed an extremely high richness of endophytes in the Xylariales, one of the largest clades of filamentous fungi and a significant source of novel secondary metabolites (SMs). Endophytes may produce host protective antimicrobial or insecticidal SMs, as well as compounds that facilitate symbiotic establishment through suppression or degradation of host immune response, but the ecological roles of most SMs are unknown. Here we characterized metabolic gene clusters in 96 genomes of endophytes and closely related saprotrophs and pathogens in two clades of Xylariales (Xylariaceae s.l. and Hypoxylaceae). Hundreds of genes appear horizontally transferred to xylarialean fungi from distantly related fungi and bacteria, including numerous genes in secondary metabolite gene clusters (SMGCs). Although all xylarialean genomes contain hyperabundant SMGCs, we show that increased gene duplications, horizontal gene transfers (HGTs), and SMGC content in Xylariaceae s.l. taxa are linked to greater phylogenetic host breadth, larger biogeographic distributions, and increased capacity for lignocellulose decomposition compared to Hypoxylaceae taxa. Overall, our results suggest that xylarialean endophytes capable of dual ecological modes (symbiotic and saprotrophic) experience greater selection to diversify SMGCs to both increase competitiveness within microbial communities and facilitate diverse symbiotic interactions.

## INTRODUCTION

Fungal endophytes inhabit asymptomatic, living photosynthetic tissues of all major lineages of plants and lichens to form one of earth’s most prevalent groups of symbionts^1^. Known from a wide range of biomes and agroecosystems^2,3^, endophytes are a ubiquitous feature of plant biology^4^. Foliar fungal endophytes are horizontally transmitted, form localized infections, and represent highly diverse and often novel lineages^3,5^. Although classified together due to ecological similar patterns of colonization, transmission, and *in planta* biodiversity^4^, endophytic fungi represent a diversity of evolutionary histories, life history strategies, and functional traits^6^.

Global, large-scale surveys of phylogenetically diverse plant and lichen hosts have revealed an extremely high richness of endophytes from boreal, temperate, tropical, and subtropical forests in the Xylariales (Sordariomycetes, Pezizomycotina, Ascomycota)^7^, one of the largest clades of filamentous fungi with >1,300 named species^8^. Previous multilocus phylogenetic examination of xylarialean endophytes in conjunction with named species (typically found as saprotrophs in decomposing leaves, wood, bark, fruits, or flowers, or more rarely as pathogens in woody hosts) demonstrated that although often closely related to named species, over half of the ∼90 endophyte taxa included in that study appeared as novel, undescribed species^7^. Moreover, the majority of xylarialean endophyte species appear to be host and substrate generalists that associate with multiple lineages of land plants and lichens, as well as in senesced leaves and leaf litter^7,9^.

In addition to their prominence as decomposers and as endophytes in a wide diversity of hosts, xylarialean fungi are a major source of novel metabolic products for use in medicine, agriculture, and industrial biofuel applications^10^. To date, >500 SMs have been described from xylarialean fungi, including various cytotoxic, antifungal, and antiparasitic agents^10^. Fungal SMs are often produced by co-localized clusters of genes that are involved in the same metabolic pathway (i.e., SM gene clusters; hereafter SMGCs)^11^. SMGCs typically contain one or more backbone genes (polyketide synthases, non-ribosomal peptide synthetases, hybrid PKS/NRPSs, terpene synthases), as well as accessory genes that modify the molecule through oxidation, reduction, methylation, or glycosylation^11^.

Xylariales genomes sequenced to date have revealed a rich repertoire of SMGCs^12^, often exceeding the numbers reported for fungi well-known for their SM production^13,14^. Such SMs may have various ecological roles: saprotrophic fungi often produce antibiotics and other toxins to inhibit microbial competitors, plant-pathogenic fungi can produce phytotoxic compounds that contribute to virulence^15^, and endophytes can produce antifungals or insect deterrents that protect their hosts^16^. Intense competition with diverse communities of soil organisms is thought to increase selection to maintain and diversify SMGCs^17^. However, we hypothesized that dual ecological modes of many xylarialean species (symbiotic and saprotrophic)^7^ may drive the horizontal gene transfers (HGTs) of SM genes and pathways necessary for both microbial competition and symbiotic establishment within diverse hosts^17–20^.

## RESULTS AND DISCUSSION

Here, we investigated the connections between fungal ecological modes and metabolic gene cluster diversity with 96 genomes within two major clades of Xylariales (Hypoxylaceae and Xylariaceae s.l., hereafter Xylariaceae), including 88 newly sequenced genomes of endophytes, saprotrophs, and plant pathogens (Fig. 1a; Supplementary Fig. 1). Taxa correspond to the previously recognized family Xylariaceae^21^ that was recently split into multiple families (Hypoxylaceae, Graphostromataceae, Barrmaeliaceae^22,23^; Supplementary Table 1). Xylarialean genomes ranged in size from 33.7-60.3 Mbp (average 43.5 Mbp; Supplementary Fig. 2b) and contained ca. 8,000-15,000 predicted genes (average 11,871; Supplementary Fig. 2c), congruent with average genome and proteome sizes of Pezizomycotina^24^. The percentage of repetitive elements per genome ranged from <1-24% (average 1.6%; Supplementary Table 2), but unlike mycorrhizal fungi^25^, repeat content was not corrected with ecological mode (Supplementary Fig. 2d).

**Figure 1.**
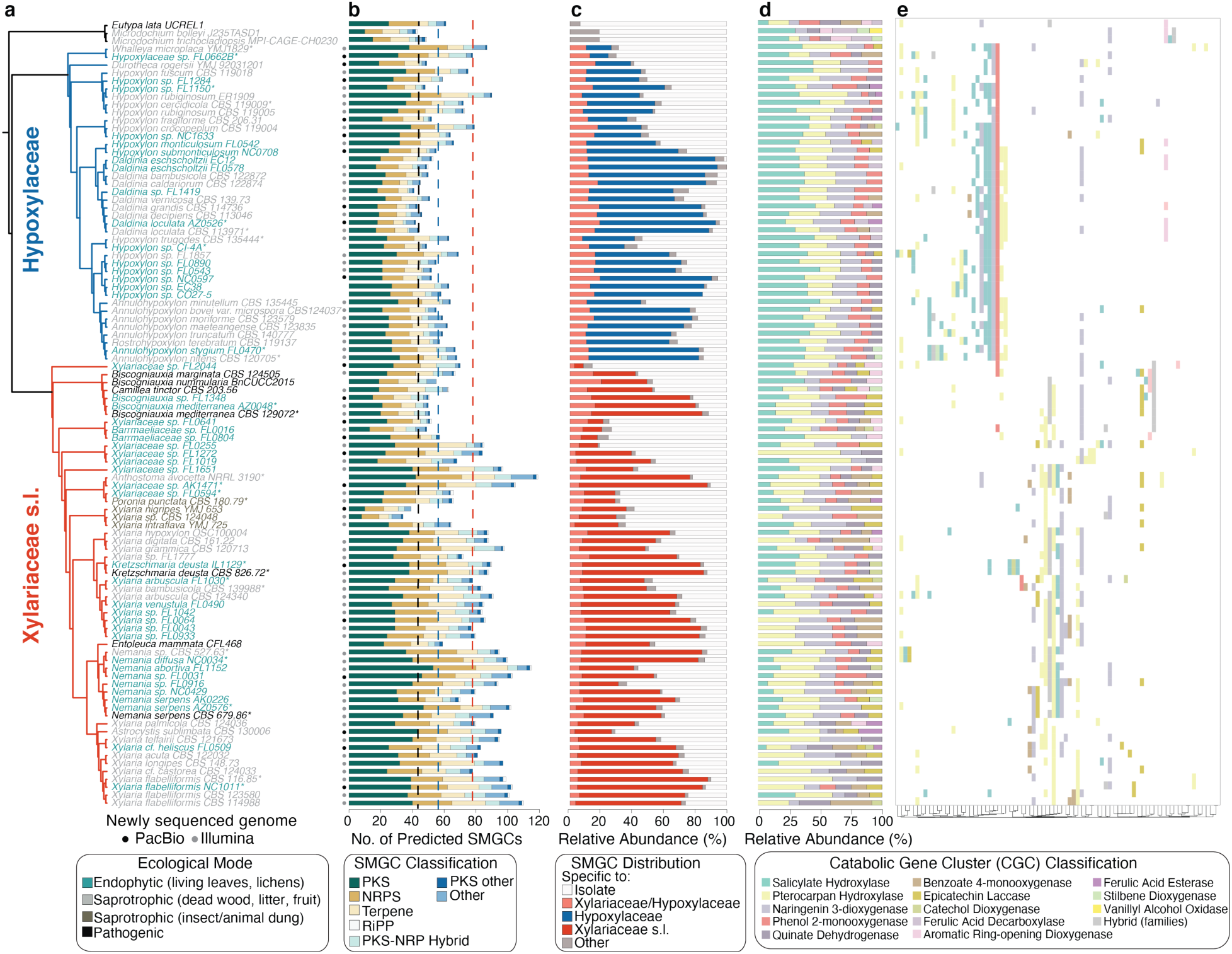
Xylariaceae s.l. and Hypoxylaceae genomes are characterized by hyperdiverse and dynamic metabolic gene clusters. (**a**) Maximum likelihood phylogenetic analyses of 1,526 universal, single-copy orthogroups support the sister relationship of the Xylariaceae s.l.^22^ (containing Xylariaceae *sensu stricto* and Graphostromataceae) and the Hypoxylaceae^23^ (Fig. 1a; Supplementary Figs. 1,2), as well as previously denoted relationships among genera^7^. Phylogenetic analyses included genomes of 25 outgroup taxa representing five other families of Xylariales and eight orders of Sordariomycetes (total 121 genomes; Supplementary Fig. 1). Taxon names are colored by ecological mode and branches colored by major clade (red: Xylariaceae s.l.; blue: Hypoxylaceae). Taxa with asterisks (*) represent 15 pairs of endophyte/non-endophyte sister taxa used to assess differences in genomic content due to ecological mode (see Fig. 5). Within this phylogenetic framework, we compared the: (**b**) abundance of different SMGC families per genome. Dotted lines indicate the averages for Pezizomycotina (black), Xylariaceae s.l. (red), and Hypoxylaceae (blue); (**c**) relative abundance of family-specific, clade-specific, and isolate-specific SMGCs; (**d**) relative abundance and (**e**) presence/absence of catabolic gene clusters (CGCs), colored by anchor gene identity (sensu^29^). Hierarchical clustering of CGCs (see bottom) was performed with the unweighted pair group method with arithmetic mean (UPGMA).

### Xylariaceae and Hypoxylaceae genomes contain hyperdiverse metabolic gene clusters

To investigate the genomic basis for the high SM production of xylarialean fungi we used antiSMASH^26^ to mine genomes for SMGCs, as well as a custom pipeline to examine metabolic gene clusters involved in the degradation of a broad array of plant phenylpropanoids^27^ (hereafter, catabolic gene clusters: CGCs). Across 96 xylarialean genomes we predicted a total of 6,879 putative SMGCs (belonging to 3,313 cluster families) and 973 putative CGCs (belonging to 190 cluster families) (Supplementary Tables 3,4). In comparison, recent large-scale analyses predicted 3,399 SMGCs (in 719 cluster families) across 101 Dothideomycetes genomes^28^ and 1,110 CGCs across 341 fungal genomes^29^. Only 25% of predicted SMGCs (n = 1,711, belonging to 816 cluster families) had BLAST hits to 168 unique MIBiG^30^ accession numbers (Supplementary Table 3b).

Total SMGCs diversity in the Xylariaceae and Hypoxylaceae is reflected in a high number of SMGCs per genome: the average number of SMGCs per genome was 71.2 (median 68), which is significantly higher than the average for fungi in the Pezizomycotina (average 42.8; Fig. 1b). At least eight xylarialean genomes contained more than 100 predicted SMGCs, with a maximum of 119 in *Anthostoma avocetta* NRRL 3190 (Fig. 1b; Supplementary Table 3). In comparison, a recent study of 24 species of *Penicillium* found an average of 54.9 SMGCs per genome, with a maximum number of 78 SMGCs observed in *P. polonicum*^13^. Genomes of Xylariaceae and Hypoxylaceae contained on average 3.3X more CGCs per genome (average 10.1; Supplementary Table 4) compared to genomes of Pezizomycotina (average 3.0^27^).

Every xylarialean genome contained SMGGs for the production of polyketides (PK; 2,871 total), non-ribosomal peptides (NRP; 2,482 total), and terpenes (1,322 total; Fig. 1b; Supplementary Table 3). SMGCs for ribosomally synthesized and post-translationally modified peptides (RiPPs) and hybrid NRP-PK compounds occurred less frequently (Fig. 1b). The most widely distributed and abundant CGCs were pterocarpan hydroxylases (n = 93), putatively involved in isoflavonoid metabolism (Fig. 1d,e; Supplementary Table 5). CGCs involved in the breakdown of plant salicylic acid^31^ (n = 251 salicylate hydroxylases) and plant flavonoids^27^ (n = 170 naringenin 3−dioxygenases) also were abundant (Fig. 1d,e). CGCs classified into nine other categories (e.g., phenol 2-monooxygenase, quinate dehydrogenase^27^) occured more rarely (Supplementary Table 4). Vanillyl alcohol oxidases, which were previously shown to be enriched in genomes of soil saprotrophs^27^, were absent in xylarialean genomes.

Consistent with the hyperdiversity of SMGCs in the Hypoxylaceae and Xylariaceae, we observed that only ca. 10% of SMGCs were shared among genomes from both Xylariaceae and Hypoxylaceae (Fig. 1c), and no SMGCs were universally present in both clades (Supplementary Table 3). On average, 21.4% and 28.2% of SMGCs per genome were unique to either taxa in the Hypoxylaceae or the Xylariaceae, respectively (range 0-82%; Fig. 1c; Supplementary Table 4), but no SMGCs were universally present within either clade. For most isolates, the majority of SMGCs were unique (i.e., ‘isolate specific’; Fig. 1c). Isolate specific SMGCs represented an average of 36.6% (SD ± 21.1) of the clusters per genome (range 0-85.7%; Fig. 1c). Even when multiple isolates of the same species were compared (e.g., *Nemania serpens* clade) 30-41% of the SMGCs appeared specific to a single isolate (Fig 1b; see also Supplementary Table 3), similar to intraspecific SMGC variation in *Aspergillus flavus*^14^.

### Impact of HGT on xylarialean genome evolution

To assess the role of HGT in shaping the genome evolution of Xylariaceae and Hypoxylaceae we performed two Alien Index (AI) analyses^32–34^. The first AI screen—designed to detect candidate HGTs from more distantly related donor lineages (e.g., bacteria, plants)—flagged 4,262 genes representing 647 orthogroups (Supplementary Table 5a). Using a custom phylogenetic pipeline (see Methods) we then identified 168 potential HGT events to Xylariaceae and Hypoxylaceae. Based on branch support and the presence of multiple xylarialean taxa in the recipient clade, 92 of these genes were deemed high-confidence HGTs (Fig. 2; Supplementary Table 5b). Similar to previous studies^35,36^, the majority of high-confidence HGTs are predicted to have been acquired from bacteria (n = 86) (Fig. 2). Other donor lineages include viruses (n = 3), Basidiomycota (n = 2) and plants (n = 1) (Fig. 2; Supplementary Table 5b). On average, xylarialean genomes had 16.2 high-confidence HGT events per genome (range: 7-30; Supplementary Table 5c). The highest number of high-confidence HGT events per genome occurred in the genome of *Xylaria flabelliformis* CBS 123580 (n = 30).

**Figure 2.**
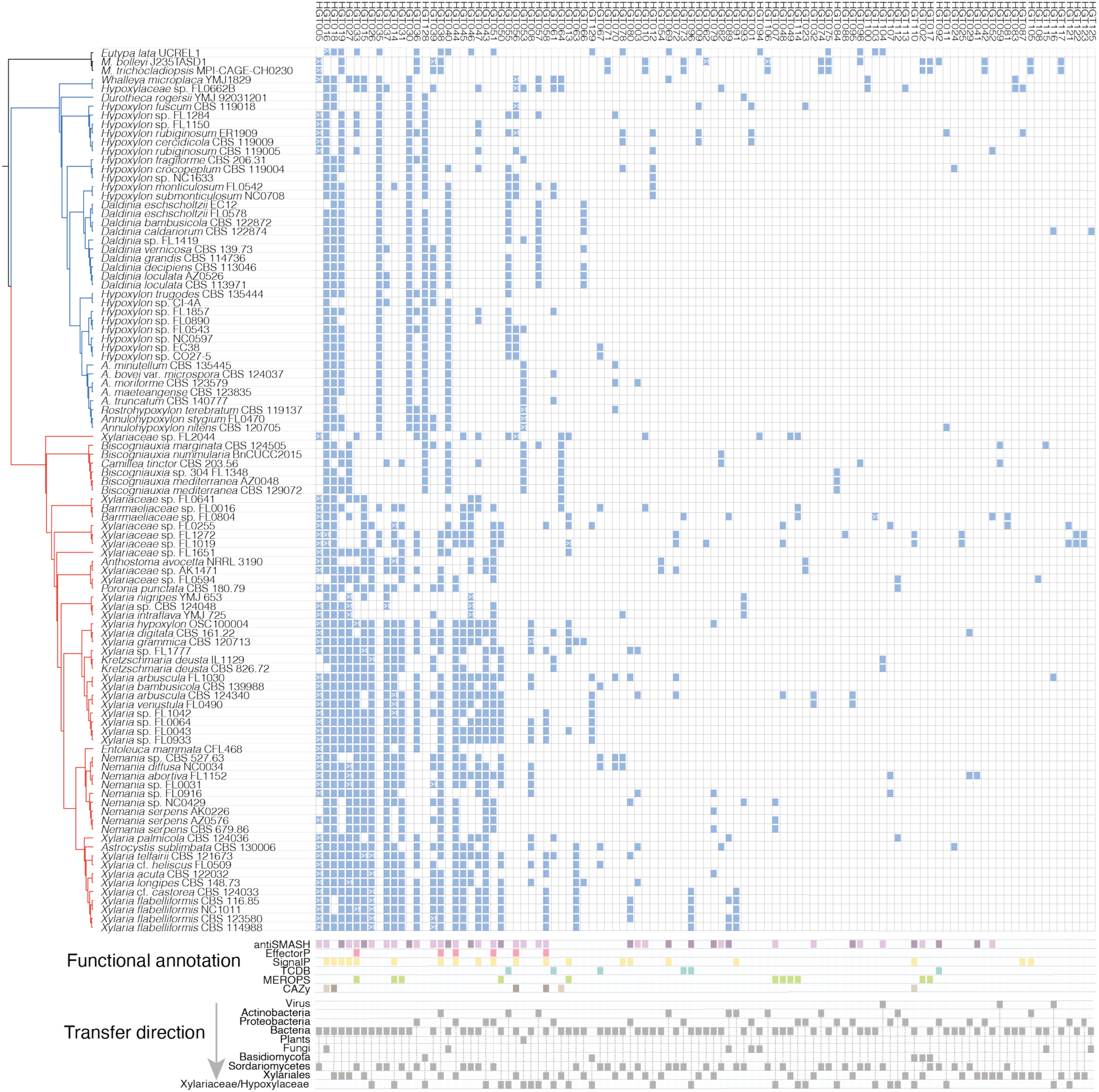
Phylogenetic distribution and functional annotation of high confidence HGTs to genomes of Xylariaceae and Hypoxylaceae. Phylogeny matches Fig. 1a. Blue boxes represent genes predicted to be high-confidence HGT events (detected with the first round of Alien Index analyses; Supplementary Table 5). HGT events are ordered from left to right based on their abundance. Transfers with more than one gene copy per genome are indicated with >1. Functional annotations (bottom) are based on antiSMASH, EffectorP, SignalP, TCDB, MEROPS, and CAZyme. SMGCs predicted as ‘biosynthetic-core’ and ‘biosynthetic-additional’ are shown with darker purple, whereas other genes in SMGCs are shown with light purple. For CAZyme predictions, dark brown color indicates plant cell wall-degrading carbohydrate-active enzyme domains (PCWDs).

HGT candidate genes were typically distributed across taxa in numerous diverse clades (n = 85 of 92 genes) rather than in monophyletic clades (Fig. 2). For example, an Enoyl-acyl carrier protein reductase protein (EC 1.3.1.9)*—*a key enzyme of the type II fatty acid synthesis (FAS) system^37^*—* occurred in bacteria (putative donor) and four distantly related recipient taxa: Xylariales sp. PMI 506, *Hypoxylon rubiginosum* ER1909; *H. cercidicola* CBS 119009; *H. fuscum* CBS 119018 (HGT0001; Supplementary Table 5). Multiple evolutionary scenarios could result in patchy taxonomic distributions. For example, multiple fungi could have independently acquired the same gene from closely related bacterial donors^36^. Alternatively, an initial HGT from bacteria to fungi may have been followed by fungal-fungal HGTs. In total, 38 HGT candidate genes occurred in genomes of both Sordariomycetes outgroup and Xylariales genomes, 28 were found in only Xylariales genomes, and 26 were only observed in genomes of Xylariaceae and Hypoxylaceae (Fig. 2; Supplementary Table 5b).

Functional annotation revealed the majority of candidate HGT genes were associated with at least one type of annotation (i.e., 95% of the highly confident and 82% of the ambiguous events; Supplementary Table 5). Six high-confidence HGT candidate genes were annotated as CAZymes, including three predicted plant cell wall degrading enzymes (PCWDEs) transferred from bacteria to diverse Xylariales (Fig. 2). No genes predicted in CGCs were identified as candidate HGTs, consistent with convergent evolution to result in similar clustering of fungal phenolic metabolism genes^27^. However, 43% of candidate HGT genes were predicted to be part of a SMGC (i.e., 40 of 92) (Fig. 2; Supplementary Tables 3,5). These include 13 genes predicted to have a biosynthetic function, such as a putative FsC-acetyl coenzyme A-N^2^-transacetylase (HGT076; Supplementary Table 5), which is part of the siderophore biosynthetic pathway in *Aspergillus* implicated in fungal virulence^38^.

Due to the high prevalence of HGT among genes predicted to be part of SMGCs, we performed a second AI screen to detect intra-fungal HGT events of genes within the boundaries of SMGCs (n = 93,066 genes) (see Methods; Supplementary Fig. 3). This analysis identified 1,148 genes in 660 SMGCs (belonging to 594 cluster families) that were putatively transferred from other fungi to members of the Xylariales (Supplementary Table 5). Candidate HGT genes were primarily for polyketide and non-ribosomal peptide production (518 PKSs, 270 NRPSs, and 180 PKS-NRPS hybrid clusters). In addition, >75% of hits to MIBiG contain genes identified by AI analyses as putative HGTs (127 of 168; see Fig. 3, bottom). SMGCs with HGT candidate genes include those with 100% similarity to MIBiG accessions from *Aspergillus*, *Fusarium*, and *Parastagonospora* involved in mycotoxin (e.g., cyclopiazonic acid, alternariol, fusarin) and antimicrobial compound (asperlactone, koraiol) production, and clusters from *Alternaria* that produce host-selective toxins (e.g., ACT-Toxin II) (Supplementary Tables 3,5). Although the second AI analysis did not identify each gene in these clusters as potential HGTs (e.g., 4 of the 19 genes in the alternariol cluster from *Hypoxylon cercidicola* CBS 119009 were predicted to be HGT; Supplementary Table 5), the phylogenetic distribution of many of these SMGCs is consistent with the acquisition of SMGCs via HGT (Fig. 3).

**Figure 3.**
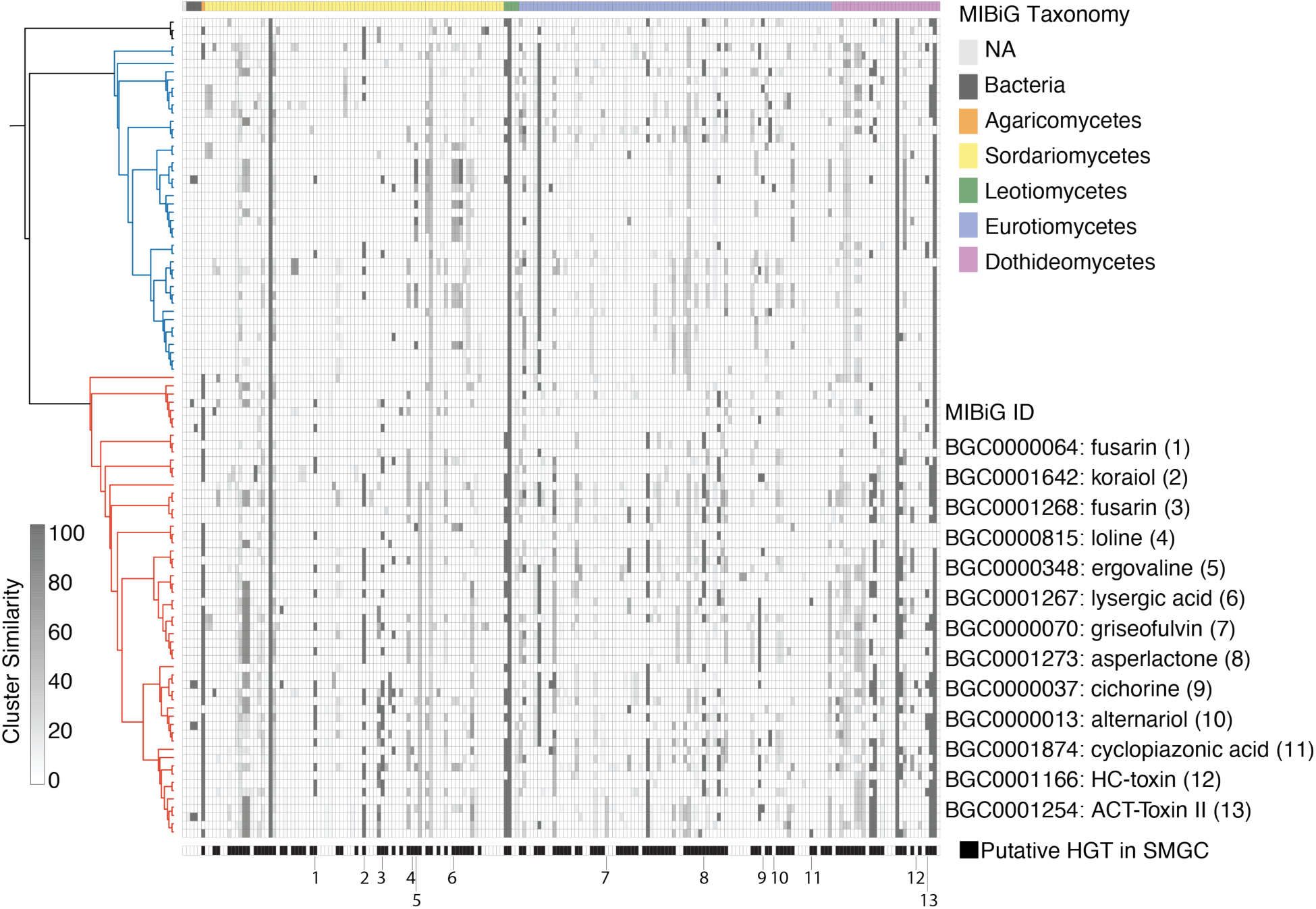
Dynamic distribution of 168 Xylariaceae and Hypoxylaceae SMGCs with hits to known metabolites in the MIBiG repository. Rows are sorted by the taxonomic identity (class and species) of the best MIBiG hit (top). Shading indicates the similarity of predicted SMGCs to reference metabolites, defined as the percentage of genes in an SMGC with significant BLAST hits to a known SMGC in the MIBiG database^39^. Black boxes (bottom) indicate SMGCs predicted by Alien Index^33,104^ to contain at least one gene putatively transferred via HGT (Supplementary Table 5). For MIBiG clusters that occurred more than once per genome, only the hit with the highest similarity is shown (Supplementary Table 3b).

Within this phylogenomic framework we also identified additional SMGCs with high similarity (i.e., calculated as the percentage of genes in an SMGC with significant BLAST hits to a known SMGC^39^) to fungal MIBiG accessions and phylogenetic distributions that support putative HGT to Xylariaceae and Hypoxylaceae (Fig. 3), but were not flagged by the second AI analysis. For example, xylarialean SMGCs with >70% similarity to clusters for ergoline alkaloids and their precursors (e.g., loline, ergovaline, and lysergic acid production) produced by Clavicipitaceae endophytes, as well as the phytotoxins cichorine cluster from *Aspergillus* (Fig. 3; Supplementary Table 3). The griseofulvin cluster from *Penicillium aethiopicum*, which produces a potent antifungal compound^40^, also appears horizontally transferred to the clade containing *X*. *castorea* and *X. flabelliformis* isolates (Fig. 3; Supplementary Fig. 4). Our analyses of HGT provide the highest support for HGTs from distantly related hosts such as bacteria (Fig. 2; see also ^36^), but our data also support fungal-fungal HGT as an important mechanism of metabolic innovation in the Xylariales. Although the discontinuous phylogenetic distributions of SMGCs observed here may represent unequal gene loss across taxa^11,17^, the presence of entire clusters known from Eurotiomycetes and Sordariomycetes in multiple endophytic and non-endophytic taxa provides additional support for HGTs.

### Expansion of Xylariaceae genomes due to increased gene duplication and HGTs

Despite the close evolutionary relationship and similar ecological niches of taxa in the Xylariaceae and Hypoxylaceae, genomes of Xylariaceae were on average ca. 7.2 Mbp larger than genomes of Hypoxylaceae (Fig. 4a; Supplementary Table 6). Larger genome size was associated with higher repeat content: Xylariaceae contained an average of 2-fold more repetitive elements (Fig. 4b; Supplementary Table 6) and had a higher density of repetitive elements surrounding genes compared to Hypoxylaceae genomes (Supplementary Fig. 5).

**Figure 4.**
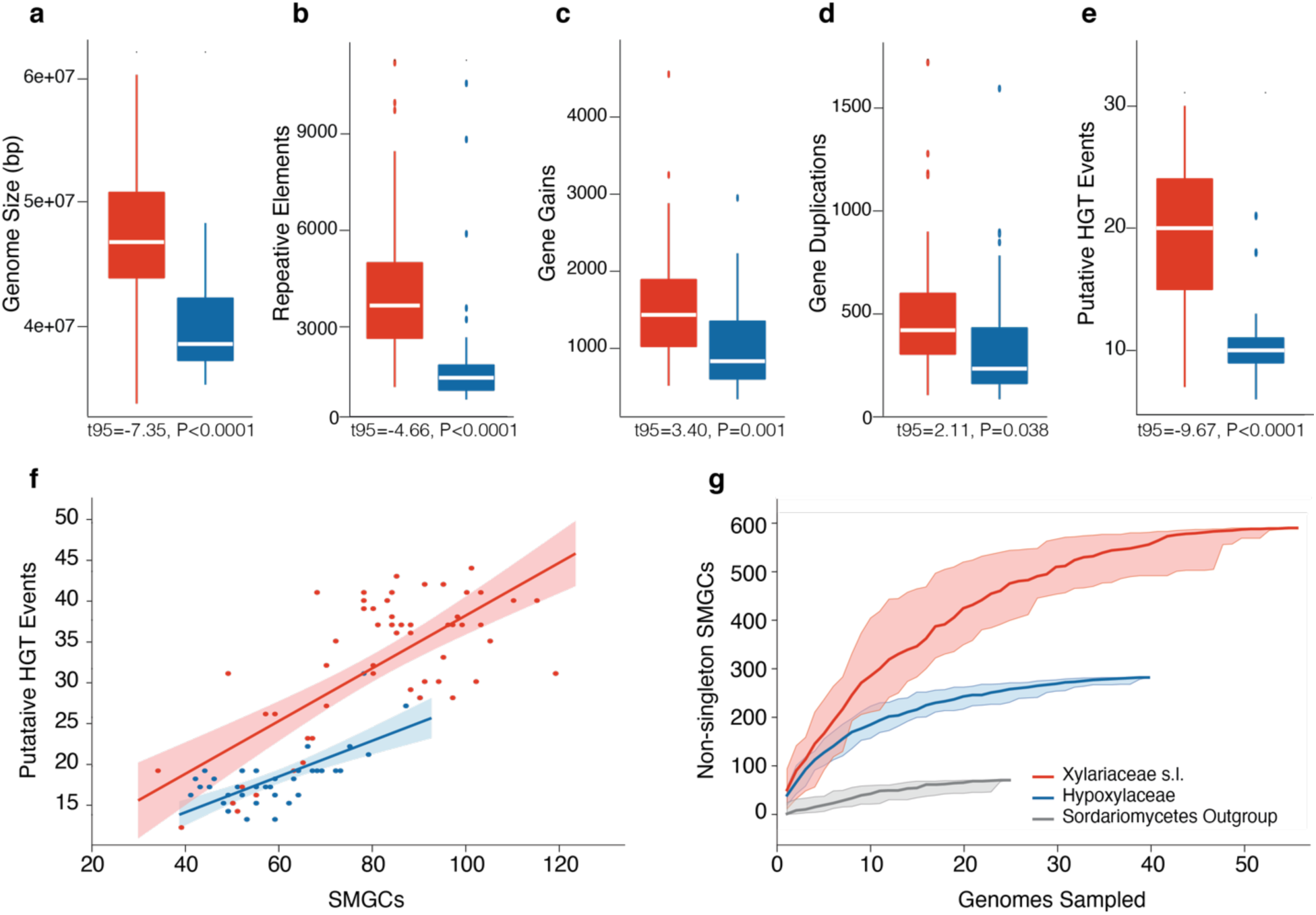
Larger genomes in the Xylariaceae clade reflect increased repetitive regions, gene gains and duplications, and HGTs. Median (**a**) genome size, (**b**) repetitive element content, (**c**) gene gains, (**d**) gene duplications, and (**e**) number of putative HGT events (high confidence only) for genomes of Xylariaceae (red) and Hypoxylaceae (blue). Box plot boundaries reflect the interquartile range. Summary statistics (averages, standard deviations, and sample sizes) are reported in Supplementary Table 6. Gene gains/losses were inferred with Wagner Parsimony under a gain penalty=loss penalty=1; (**f**) Relationship between the number of HGT events and SMGCs as a function of clade (Pearson correlation for each clade was the same; r = 0.72, P<0.0001); (**g**) Rarefaction curves of non-singleton SMGCs by clade illustrates that when compared at the same number of genomes (n = 25), the richness of SMGCs is highest in the Xylariaceae. SMGC richness for Xylariaceae and Hypoxylaceae genomes is ca. 4-7X greater than outgroup genomes (n = 71 SMGCs).

In addition to greater repeat content, Xylariaceae genomes also contained on average 750 more protein-coding genes compared to Hypoxylaceae genomes (P<0.0001; Supplementary Table 6b). Ancestral state reconstructions reveal that Xylariaceae genomes have experienced significantly more gene gains (n = 472), gene duplication events (n = 136), orthogroup gains (n = 313), and orthogroup expansion events (n = 90) compared to Hypoxylaceae clade since the radiation from their last common ancestor (Fig. 4c-d), although both clades underwent similar numbers of gene losses (t_95_ = 0.51, P=0.61; Supplementary Table 6b). Xylariaceae genomes also experienced on average ca. 2-fold more HGTs events compared to Hypoxylaceae genomes (Fig. 4e).

Increased genome sizes resulting from HGT were positively associated with increased numbers of SMGCs across both clades (Fig. 4f), reflecting the fact that clustered metabolite genes in fungi are more likely to undergo HGT compared to unclustered genes^41^. Genomes of Xylariaceae contained on average ca. 20 more SMGCs than Hypoxylaceae genomes (Supplementary Table 6b) and ca. 2-fold greater cumulative richness of SMGCs compared to Hypoxylaceae clade (2,336 vs 1,075 total; 587 vs 282 non-singleton). Rarefaction analysis reveals the richness of non-singleton SMGCs increases at a greater rate in the Xylariaceae clade (Fig. 4g). Genomes of Xylariaceae also contained a greater fraction of isolate specific SMGCs compared to Hypoxylaceae, regardless of SMGC type (Xylariaceae: 31.2 ± 16.1; Hypoxylaceae: 19.8 ± 15.3; P = 0.0007; Fig. 1c; Supplementary Fig. 6). Yet despite the high variation of SMGCs among taxa, network analysis illustrates that the composition of SMGCs is more similar among isolates from the same clade, regardless of ecological mode (Supplementary Fig. 7).

In contrast to the pattern observed for SMGCs, genomes of Hypoxylaceae contained a greater number of CGCs than Xylariaceae genomes (Xylariaceae: 9.5 ± 0.4; Hypoxylaceae: 11.0 ± 0.4; P = 0.0068; Supplementary Table 4) and different classes of CGC dominated the two clades (Fig. 1d,e). For example, salicylate hydroxylases were the most abundant CGCs among Hypoxylaceae, but were absent from 25% Xylariaceae genomes (Fig. 1d). In contrast, CGCs classified as pterocarpan hydroxylases were the most abundant CGC type in genomes of Xylariaceae (Fig. 1d). Four types of CGCs were universally present across Hypoxylaceae: salicylate hydroxylase, pterocarpan hydroxylase, naringenin 3-dioxygenase, phenol 2-monooxygenase (Fig. 1d). CGCs classified as naringenin 3-dioxygenase were the only CGC type found across all Xylariaceae genomes.

In addition to different metabolic gene clusters content, additional differences between Xylariaceae and Hypoxylaceae genomes suggest different functional capacities. Xylariaceae genomes contain greater numbers of genes with signaling peptides, as well as genes annotated as effectors, membrane transport proteins, transcription factors, peptidases, and CAZymes compared to Hypoxylaceae, even after accounting for differences in genome size (Supplementary Table 6). For example, on average genomes of Xylariaceae contained ca. 50 more CAZymes than Hypoxylaceae (Xylariaceae 579.9 ± 7.7; Hypoxylaceae 529.6 ± 9.1, P <0.0001), including a significant increase in PCWDEs involved in the degradation of cellulose, hemicellulose, lignin, pectin, and starch (Supplementary Table 6). Additionally, comparison of gene ontology (GO) terms for shared orthogroups significantly enriched in either Xylariaceae or Hypoxylaceae (i.e., 74 and 26, respectively) revealed that the Hypoxylaceae had a significant increase in the number of GO terms associated with membrane transport, whereas Xylariaceae had a significant increase in the number of GO terms for catalytic activities and binding (Supplementary Fig. 8b).

### Xylariaceae genome evolution linked to ecological generalism

The majority of described Xylariaceae and Hypoxylaceae species are wood- or litter-degrading saprotrophs or woody pathogens^42,43^, although both culture-based and culture-free studies of healthy photosynthetic tissues of plants and lichens demonstrate the abundance and novel diversity represented by xylarialean endophytes^7^. Previous studies have identified isolates with highly similar ITS nrDNA sequences occurring in both living host tissues as well as decomposing plant materials^7,9^, which suggests that endophytism may represent only part of a complex life cycle that blurs the lines between distinct ecological modes^7^.

In support of such ecological generalism, we observed no clear distinctions in genome size or content among endophytic and non-endophytic taxa when all ingroup genomes were analyzed (Supplementary Table 6). One exception was the reduced genomes and CAZyme content of termite-associated *Xylaria* spp. (i.e., *X. nigripes* YMJ 653, *X.* sp. CBS 124048, and *X. intraflava* YMJ725; Supplementary Fig. 2a, Supplementary Fig. 9) that reflects a single evolutionary transition to specialization on termite nest substrates decomposed by a basidiomycete fungus^43^. The lack of clear genomic signal for endophytism in the Xylariaceae and Hypoxylaceae contrasts sharply with genome evolution in ectomycorrhizal fungi, where mycorrhizal clades have experienced convergent loss of genes that encode lignocellulose-degrading enzymes and an increase in small secreted effector-like proteins since their divergence from saprotrophic ancestors^25^. However, a recent analysis of 101 ecologically diverse Dothideomycetes revealed only six orthogroups predicted plant-pathogenic vs. saprotrophic ecological mode with >95% accuracy^44^, highlighting the complexity of linking genotype to phenotype for complex traits.

Despite their ecological similarities, genomes of Xylariaceae experienced more gene duplications, gene family expansions, and HGT events, resulting in higher SMGC content as well as more genes important for pathogenicity (e.g., effectors, peptidases) and saprotrophy (e.g, CAZymes, transporters) in comparison with Hypoxylaceae (Supplementary Table 6). As genomes of fungi with saprotrophic lifestyles typically contain more CAZymes and PCWDEs compared to plant pathogens and mycorrhizal symbionts^25,44,45^, our genomic results are consistent with the potential for Xylariaceae fungi (including endophytes) to have greater saprotrophic abilities compared to Hypoxylaceae fungi^46^. To test this prediction, we compared the abilities of 20 isolates to degrade leaves of *Pinus* and *Quercus*. We found that isolates of Xylariaceae with expanded CAZymes and PCWDEs repertoires caused greater mass loss compared to taxa with fewer genes predicted to degrade lignocellulose (i.e., Hypoxylaceae and Xylariaceae from animal-dung clade; Supplementary Fig. 10).

The genomic and functional differences we observed are consistent with Xylariaceae species as ecological generalists encompassing both endophytic and saprotrophic life stages. Xylariaceae endophyte species also associate with a greater phylogenetic diversity of plant and lichen hosts compared to species of Hypoxylaceae endophytes (t_42_ = 2.25; P = 0.0294; Supplementary Fig. 11a). Host breadth of Xylariaceae endophytes also is positively associated with the number of total HGT events and the number of SMGCs for non-ribosomal peptides (Supplementary Fig. 11b). Thus, we hypothesized Hypoxylaceae taxa may have undergone fewer HGT events and gene expansions due to species having more distinct ecological modes with less selection for metabolic versatility. To test this hypothesis, we performed pairwise comparisons of 15 sister taxa across both clades with contrasting ecological modes, which revealed that endophyte genomes in the Hypoxylaceae contain significantly fewer genes with signaling peptides, protein coding genes, transporters, peptidases, PCWDEs (especially those involved in decomposition of cellulose and lignin), SMGCs, and CGCs than non-endophyte genomes (Fig. 5). In contrast, no significant differences were observed between endophytes and saprotrophs in the Xylariaceae clade (Fig. 5; Supplementary Table 6).

**Figure 5.**
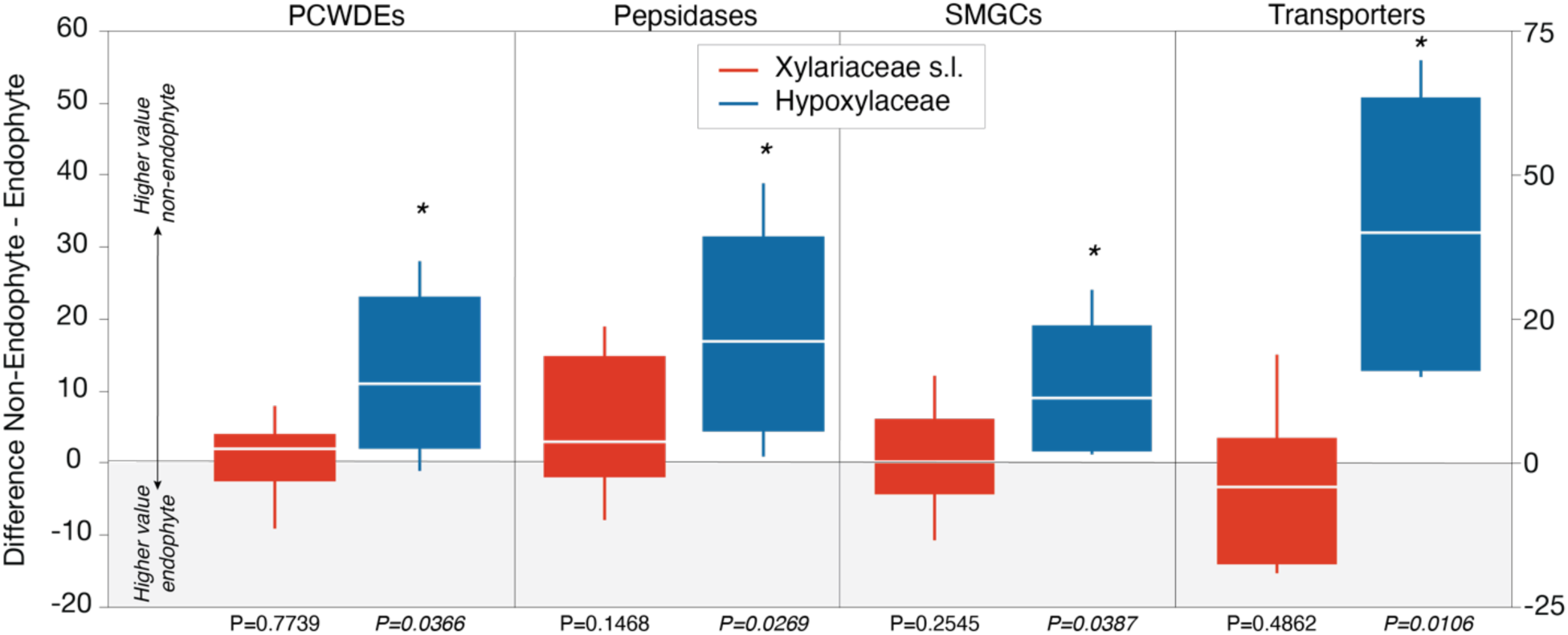
Pairwise comparisons of sister taxa illustrate ecological modes are more distinct in the Hypoxylaceae. Box plots of the median and interquartile difference in gene counts of PCWDEs, peptidases, SMGCs (y-axis on left), and transporters (y-axis on right) between 15 pairs of sister taxa with contrasting ecological modes for Xylariaceae and Hypoxylaceae (sister taxa are indicated with asterisks in Fig. 1a). Values greater than zero indicate higher gene counts in non-endophytic taxa, whereas differences less than zero indicate higher gene counts in endophytes. Statistical differences were assessed with least squares means contrast under the null hypothesis: non-endophyte value -endophyte value = 0 (see Supplementary Table 6 for summary statistics). P-values <0.05 are indicated with an asterisk (*).

Increased metabolic diversity and host breadth of Xylariaceae species likely impacts their geographical distributions^47^. Xylariaceae genera such as *Xylaria* and *Nemania* occur worldwide as fruiting bodies in temperate, subtropical, and tropical forests, whereas Hypoxylaceae genera such as *Daldinia* and *Hypoxylon* are more common in boreal and temperate forests^7^, but taxonomic uncertainty for many specimens and sequences^48^ combined with a lack of biome metadata for the majority of reference taxa^7^ precludes robust statistical comparisons of biogeographic ranges for named taxa. However, a recent global survey of boreal endophytes demonstrated that host generalist species occupy larger geographic ranges^3^ and re-analysis of data from previous ecological surveys in boreal, temperate montane, and subtropical forests in Alaska, Arizona, North Carolina, and Florida reveals a higher fraction of Xylariaceae endophyte species cultured from hosts in more than one site (i.e., 28% Xylariaceae vs. 20% for Hypoxylaceae), including six Xylariaceae endophyte species that were found in >3 sites^7^. In contrast, no Hypoxylaceae endophyte species from that study were found in more than two sites^7^.

## CONCLUSIONS

Our analysis of 96 phylogenetically and ecologically diverse Xylariaceae and Hypoxylaceae genomes reveals that gene duplication, gene family expansion, and HGT of SMGCs from putative bacterial and fungal donors, drives metabolic versatility in the Xylariaceae. Expanded metabolic diversity of Xylariaceae taxa is associated with greater phylogenetic host breadth, larger biogeographic distributions, and increased capacity for lignocellulose decomposition compared to Hypoxylaceae taxa. Yet despite differences among clades, our data suggest that saprotrophs in both clades are under selection to maintain both large gene repertoires to degrade diverse lignocellulosic compounds^44^ and highly diverse SMGCs that likely increase competitive abilities in diverse microbial communities^11,49,50^ (Supplementary Table 6e). In contrast, SMGC abundance in endophyte genomes appears unrelated to PCWDE content. Overall, our results provide evidence that SMGCs may play a key role in facilitating endophyte colonization of diverse hosts (e.g., through suppression or degradation of host immune responses^17,18^), further highlighting the importance of symbioses to drive not only speciation and ecological diversification^51^, but chemical biodiversity that can be leveraged for novel pharmaceuticals and agrochemicals^52^.

## METHODS

### Fungal strain selection and verification

Isolates were selected based on their phylogenetic position (previously estimated using multilocus phylogenetic analyses^7^), as well as their ecological mode (i.e., endophyte, saprotroph, pathogen). To minimize the effect of phylogeny when assessing the impact of ecological mode on genome evolution, we included 15 pairs of closely related sister taxa with contrasting ecological modes (i.e., endophyte vs. non-endophyte)^7^. Ecological modes were assigned based on the substrate of isolation: fungi isolated from living plants and lichens with no signs of disease were classified as endophytic; fungi isolated from or collected as fruiting bodies from decomposing plant tissues (e.g., litter, wood, dung) were classified as saprotrophs; and fungi isolated or collected as fruiting bodies from living, diseased host tissues were classified as pathogens. For strains that lacked host and substrate metadata, ecological modes were estimated based on information for that species in the literature (see ref ^7^).

In total, we sequenced 44 endophytic taxa^2,53^ and 44 named taxa of Xylariaceae s.l. and Hypoxylaceae (Supplementary Table 1). Endophytic isolates are maintained as an axenic voucher in sterile water at the Robert L. Gilbertson Mycological Herbarium at the University of Arizona (ARIZ). Cultures of named taxa were obtained from the Westerdijk Fungal Biodiversity Institute (Netherlands) or from Dr. Yu-Ming Ju. In total, we sequenced genomes representing ca. 24 genera and 80 species of Xylariaceae s.l. and Hypoxylaceae, as well as an additional two undescribed species of endophytic Xylariales (*Pestalotiopsis* sp. NC0098 and Xylariales sp. AK1849) included in the outgroup (Supplementary Fig. 1).

Prior to genome and transcriptome sequencing, fungi were grown on 2% malt extract agar (MEA) to verify morphology and obtain tissue for a preliminary DNA extraction to verify isolate identity. Briefly, DNA was extracted using Extract n Amp (Sigma) following ref ^54^. For each isolate the ITS-LSU nrDNA region was PCR amplified using the primer pair ITS1F/LR3 and Sanger sequenced for each isolate as described by ref ^2^. Sequences were edited in Sequencher v5.4.6 (Gene Codes Corporation, Ann Arbor, MI) and aligned with the original ITS nrDNA sequences for each isolate. For isolates without a prior ITS nrDNA sequence, we used T-BAS v2^55^ to query sequences against the multilocus tree of the Xylariaceae from ref ^7^. In some cases, names of reference taxa (previously named based only on morphological characters) were updated to reflect their phylogenetic placement (see Supplementary Table 1).

### DNA and RNA purification

After strains were verified, we used two different mycelial growth and cultivation techniques to achieve the specific nucleic acid concentration and quality requirement for either Illumina or PacBio Single-Molecule Real-Time (SMRT) sequencing. For PacBio sequencing, isolates were first grown on multiple 2% MEA plates overlaid with sterile, cellophane membrane to allow mycelial harvesting without media carry-over. After ca. 5-10 days of growth, mycelium was removed using sterile forceps and scalpels, placed in 150 mL of 1% malt extract (ME) media in a sterile, stainless steel Eberbach blender cup (Fisher Scientific) and homogenized with 3-5 short pulses using a Waring blender. After homogenization, two 75 mL aliquots were placed in Erlenmeyer flasks and incubated on a shaker at room temperature for 3-7 days. Once sufficient growth was obtained samples were then filtered through sterilized Miracloth (Millipore, 475855-1R) in a Buchner funnel (Fisher Scientific), placed in a 50 mL centrifuge tube, flash-frozen in liquid nitrogen, and stored at -80°C. If isolates grew slowly, the contents of the inoculated flask were re-blended with an equal volume of fresh 1% ME media after 7 days, aliquoted into new flasks, and incubated on the shaker at room temperature for an additional 5-7 days prior to filtering. After filtration, mycelium was washed with sterile molecular grade water to remove media and excess polysaccharides.

DNA isolation for PacBio sequencing was performed using a modified phenol:chloroform extraction method (see ref ^56^). Briefly, ca. 4 g (wet weight) of tissue was ground in liquid nitrogen with a sterile mortar and pestle. Ground tissue was transferred to a 50 mL Falcon tube containing 14 mL of SDS buffer and incubated at 65°C for 30 minutes, during which the tube was gently inverted 5X every 10 minutes. After incubation, 0.5X volume of 5M KOAc (pH 7.5) was added to each tube, mixed by inversion, and placed at 4°C for 30 minutes. Samples were then centrifuged at 4500 RPM for 10 minutes at 4°C. After centrifugation, the supernatant was removed, placed into a new tube, 0.7X volume of molecular grade isopropanol was added, and the tube was gently inverted to mix. The sample was then centrifuged at 4500 RPM for 20 minutes at 4°C to precipitate the DNA. After centrifugation the supernatant was removed, and the DNA pellet was washed with 5 mL of 70% EtOH and centrifuged for an additional 5 minutes at 4500 RPM. Residual EtOH was removed with a pipette, and the pellet was air dried. The DNA pellet was resuspended in 2 mL of TE buffer, 10 uL of RNase (20mg/mL; Invitrogen, Waltham, MA) and the sample was placed in a 37°C water bath for 1 hour. After incubation, DNA was purified with phenol:chloroform:IAA, washed with 0.3X volume of absolute molecular grade ethanol to remove polysaccharides, and precipitated by adding 1.7X volume of absolute molecular grade ethanol. The resulting DNA pellet was washed with 70% EtOH, air dried, and resuspended in low salt TE.

For Illumina sequencing, isolates were first grown on multiple 2% MEA plates overlaid with sterile cellophane as described above, but harvested mycelium was placed in RNase free stainless-steel bead tubes (Next Advance, NAVYR5-RNA), flash frozen in liquid nitrogen, and stored at -80°C until extraction. DNA for Illumina sequencing was extracted using similar methods as above for PacBio, with the exception that only a small amount of tissue was used, samples were homogenized in 2 mL tubes with stainless steel beads rather than grinding in liquid N, and the initial purification with 5M KOAc was not performed (see ref ^57^). DNA obtained from both methods was quantified with a Qubit fluorometer (Invitrogen, Carlsbad, CA) and sample purity was assessed with a NanoDrop 1000 (BioNordika, Herlev, Denmark). The purity of DNA for PacBio sequencing was also verified with a EcoRI (New England BioLabs, Ipswich, MA) restriction digest and sized via electrophoresis on a 1% agarose gel with a clamped homogeneous electric field (CHEF) apparatus^58^ as described in ref ^59^.

RNA was extracted for each isolate with the Ambion Purelink RNA Kit (Thermo Fisher Scientific, Waltham, MA). Briefly, isolates were grown on 2% MEA with sterile cellophane overlay. Mycelium was harvested after ca. one week of growth, placed in 2 mL tubes containing stainless steel beads, flash frozen in liquid N, and stored at -80°C until extraction. Frozen mycelium was homogenized for 5 seconds at 1400 RPM on a BioSpec, Mini-BeadBeater 96 115V (MP Biomedicals) with stainless steel beads. Following homogenization, 1 mL of TRIzol was added to each tube and the sample was incubated for 5 minutes at room temperature, followed by centrifugation at 4°C for 15 minutes at 12,000 RPM. Following centrifugation, the supernatant was transferred to a new tube and 0.2 mL of chloroform was added, mixed gently by inversion, and transferred to a column following the manufacturer’s instructions. RNA was quantified with a Qubit fluorometer (Invitrogen) and sample purity was assessed with a NanoDrop (BioNordika). RNA was then treated with DNase (Thermo Fisher Scientific) following the manufacturer’s instructions and RNA integrity was assessed on a BioAnalyzer at the University of Arizona Genomics Core Facility.

### Genome and transcriptome sequencing and assembly

Genomes were generated at the Department of Energy (DOE) Joint Genome Institute (JGI) using Illumina and PacBio technologies (Supplementary Table 1). For 66 isolates, Illumina standard shotgun libraries (insert sizes of 300bp or 600bp) were constructed and sequenced using the NovaSeq platform. Raw reads were filtered for artifact/process contamination using the JGI QC pipeline. An assembly of the target genome was generated using the resulting non-organelle reads with SPAdes^60^. PacBio SMRT sequencing was performed for 22 isolates of Xylariaceae s.l. and Hypoxylaceae and two additional endophytic Xylariales (Xylariales spp. NC0098 and AK1849) on a PacBio Sequel. Library preparation was performed either using the PacBio Low Input 10kb or PacBio >10kb with AMPure Bead Size Selection. Filtered sub-read data were processed with the JGI QC pipeline and *de novo* assembled using Falcon (SEQUEL) or Flye (SEQUEL II). Stranded RNASeq libraries were created and quantified by qPCR. Transcriptome sequencing was performed on an Illumina NovaSeq S4. Raw reads were filtered and trimmed using the JGI QC Pipeline.

Plate-based RNA sample prep was performed on the PerkinElmer Sciclone NGS robotic liquid handling system using Illumina’s TruSeq Stranded mRNA HT sample prep kit utilizing poly-A selection of mRNA following the protocol outlined by Illumina in their user guide (https://support.illumina.com/sequencing/sequencing_kits/truseq-stranded-mrna.html) and with the following conditions: 1 ug of total RNA per sample and eight cycles of PCR for library amplification. The prepared libraries were quantified using KAPA Biosystems’ next-generation sequencing library qPCR kit and run on a Roche LightCycler 480 real-time PCR instrument. Sequencing of the flowcell was performed on the Illumina NovaSeq sequencer using NovaSeq XP V1 reagent kits, S4 flowcell, following a 2×150 indexed run recipe. Raw reads were evaluated with BBDuk (https://sourceforge.net/projects/bbmap/) for artifact sequences by kmer matching (kmer=25), allowing 1 mismatch and detected artifacts were trimmed from the 3’ end of the reads. RNA spike-in reads, PhiX reads, and reads containing any Ns were removed. Quality trimming was performed using the phred trimming method set at Q6. Following trimming, reads under the length threshold were removed (minimum length 25 bases or 1/3 of the original read length - whichever is longer). Filtered reads were assembled into consensus sequences using Trinity v2.3.2^61^ with the -- normalize_reads (In-silico normalization routine) and --jaccard_clip (Minimizing fusion transcripts derived from gene dense genomes) options.

### Genome annotation

Genomes were annotated using the JGI annotation pipeline^62^. Functional annotations were obtained from InterPro^63^, PFAM^64^, Gene Ontology (GO^65^), Kyoto Encyclopedia of Genes and Genomes (KEGG^66^), Eukaryotic Orthologous Groups of Proteins (KOG^67^), the Carbohydrate-Active EnZymes database (CAZy^68^), MEROPS database^69^, the Transporter Classification Database (TCDB^70^), and SignalP v3.0a^71^. CAZymes involved in the degradation of the plant cell wall were separated according to ref ^72^. Annotation information for each isolate is available through MycoCosm^62^. We examined repetitive elements using RepeatScout^73^, which identifies novel repeats in the genomes, and RepeatMasker (http://repeatmasker.org), which identifies known repeats based on the Repbase library^74^. Candidate effectors were predicted using EffectorP v2.0^75^. Genome sequencing yielded eukaryotic Benchmarking Universal Single-Copy Orthologs (BUSCO) values ≥95% (Supplementary Table 1). On average, ca. 90% of RNAseq reads mapped to each genome (Supplementary Table 1).

### Orthogroup prediction

For comparative analyses, data for 23 additional genomes of Sordariomycetes were obtained from MycoCosm^62^, including outgroup taxa belonging to the Hypocreales (n = 6)^76–81^, Glomerellales (n = 3)^82–84^, Sordariales (n = 2)^85^, Coniochaetales (n = 1)^86^, Togniniales (n = 1)^87^, Diaporthales (n = 1)^88^, Magnaporthales (n = 1)^89^, Ophiostomatales (n = 1)^90^, and five other families of Xylariales (n = 7)^91–93^. Data from an additional eight taxa in Xylariaceae *sensu lato*^94^ also were obtained from MycoCosm^62^ (Supplementary Fig. 1; Supplementary Table 1). Orthologous gene families (i.e., orthogroups) for all 121 genomes (ingroup and outgroup) were inferred by OrthoFinder v2.3.3^95^, which was executed using DIAMOND v0.9.22^96^ for the all-versus-all sequence similarity search and MAFFT v7.427^97^ for sequence alignment.

### Phylogenomic analysis

Protein sequences of 1,526 single-copy orthogroups defined by OrthoFinder were aligned using MAFFT v7.427^97^, concatenated, and analyzed using maximum-likelihood in IQ-TREE multicore v1.6.11^98^ with the Le Gascuel (LG) substitution model. Node support was calculated with 1,000 ultrafast bootstrap replicates. Additional phylogenomic analyses with different models of evolution, gene sets, and outgroup taxa resulted in nearly identical topologies (see Supplementary Materials; Supplementary Fig. 12).

### Analysis and functional annotation of orthologous gene families

Representative annotations from InterPro^63^, PFAM^64^, GO^65^, CAZy^68^, MEROPS^69^ and TCDB^70^, SignalP 3.0^71^, and EffectorP 2.0^75^ were assigned to orthogroups KinFin v1.0^99^. The criteria for orthogroup annotation was (i) a minimum of 75% of the proteins in the orthogroup share the annotation and (ii) 30% of the taxa in the cluster with at least one protein annotated with that domain. KinFin was used to aid in the classification of orthogroups into different categories such as isolate-specific, subfamily-specific, and universal (Supplementary Fig. 2e; Supplementary Table 7) and to identify orthogroups that were significantly enriched or depleted in the two major clades (i.e., Xylariaceae and Hypoxylaceae) using the Mann-Whitney U test (Supplementary Fig. 8a). We also compared functional categories for universal (i.e,. “core”) vs. isolate-specific (i.e.,“dispensable”) orthogroups using euKaryotic Orthologous Groups (KOGs) (Supplementary Fig. 13; Supplementary Table 3f). We also used KinFin to generate a network representation of the OrthoFinder clustering. The resulting network was edited with Gephi v0.9.1^100^, whereby nodes were positioned by a force directed layout algorithm (as described by ref ^99^) (Supplementary Fig. 14).

### Ancestral state reconstruction of orthologous gene family size

We used ancestral gene content reconstruction in Count v10.04 ^101^ with the unweighted Wagner parsimony method (gain and loss penalties both set to 1) to assess changes in the size of orthologous gene families over evolutionary time. This gene tree unaware method requires as input the organismal phylogeny and a gene family size table showing the numbers of genes per orthogroup per taxa, estimates the gene family sizes, as well as orthogroup gain and loss events at ancestral nodes. An orthogroup gain is defined as a shift from orthogroup absence at the preceding node to presence at the node of interest, and orthogroup loss is the opposite transition. Functional annotation of orthogroups was imported into Count v10.04 GUI to assist in the interpretation of the results. The ancestral gene content was reconstructed for the entire data set, as well as for subsets of orthologous gene families corresponding to CAZymes and PCWDEs (see Supplementary Fig. 15).

### Metabolic gene cluster prediction

SMGCs were predicted using antiSMASH version 5.1.0^26^ setting the strictness to ‘relaxed’ and enabling ‘KnownClusterBlast’, ‘ClusterBlast’, ‘SubClusterBlast’, ‘ActiveSiteFinder’, ‘Cluster Pfam analysis’ and ‘Pfam-based GO term annotation’. Clinker and clustermap.js were used to visualize and compare SMGCs^102^. Sequence similarity network analysis of the SMGCs was performed using the Biosynthetic Gene Similarity Clustering and Prospecting Engine (BiG-SCAPE) v1.0.1^103^. BiG-SCAPE was executed under the hybrid mode, enabling the inclusion of singletons and the SMGCs from the Minimum Information about a Biosynthetic Gene cluster (MIBiG) repository version 1.4^30^. To compare the distribution of SMGCs, BiG-SCAPE families representing different SMGC types were combined into a single dataset. To remove duplicates, SMGCs assigned to multiple families were arbitrarily assigned to the largest family. The output from BiG-SCAPE was also incorporated into KinFin^99^ to visualize gene content similarity as network graphs (Fig. 2d) as well as examine SMGC distribution across clades (Fig. 2b). We observed no correlation of SMGC content and the number of scaffolds per genome (Supplementary Fig. 16).

To examine metabolic gene clusters involved in catabolism (i.e., catabolic gene clusters; CGCs) we used cluster_retrieve (https://github.com/egluckthaler/cluster_retrieve) to search for clusters containing phenylpropanoid degradation “anchor” genes^27^. Cluster_retrieve searches for multiple “cluster models” containing one of 13 anchor genes: aromatic ring-opening dioxygenase (ard), benzoate 4-monooxygenase (bph), ferulic acid esterase 7 (cae), catechol dioxygenase (cch), epicatechin laccase (ecl), ferulic acid decarboxylase (fad), pterocarpan hydroxylase (mak), naringenin 3-dioxygenase (nad), phenol 2-monooxygenase (pmo), quinate 5-dehydrogenase (qdh), salicylate hydroxylase (sah), stilbene dioxygenase (sdo), and vanillyl alcohol oxidase (vao)^27^. Homologous genes in each locus were defined by a minimum BLASTp (v2.2.25+) bitscore of 50 and 30% amino acid identity, and target sequence alignment 50-150% of the query sequence length. Homologs of query genes were considered clustered if separated by no more than six intervening genes. Clusters on the same contig were consolidated if separated by less than 30kb and homologous cluster families across genomes were inferred using a modified version of BiG-SCAPE^103^. We adapted the BiG-SCAPE network model for catabolic clusters by adding catabolic anchor genes to “anchor_domains.txt” and manually tuning the “Others” cluster type model parameters until known related clusters, such as quinate dehydrogenase clusters, merged into families. Tuning resulted in the values 0.35 for the Jaccard dissimilarity of cluster Pfams, 0.63 for Pfam sequence similarity, 0.02 adjacency index, and 2.0 anchor boost.

### Detection of HGT events

We used the Alien Index (AI) pipeline (https://github.itap.purdue.edu/jwisecav/wise) as previously described (see refs ^33,104^) to identify HGT candidates across the 121 genomes. Each predicted protein sequence was queried against a custom protein database using Diamond v0.9.22.123^96^. The custom database consisted of NCBI RefSeq (release 98)^105^ supplemented with additional predicted protein sequences from the Marine Microbial Eukaryotic Transcriptome Sequencing Project (MMETSP)^106^ and the 1000 Plants transcriptome sequencing project (OneKP)^107^. Diamond results were sorted based on the normalized bitscore (*nbs*), where *nbs* was calculated as the bitscore of the single best high scoring segment pair (HSP) in the hit sequence divided by the best bitscore possible for the query sequence (i.e., the bitscore of the query aligned to itself).

To identify HGT candidates, an ancestral lineage is first specified, and the AI score calculated using the formula: AI=*nbsO*-*nbsA*, where *nbsO* is the normalized bit score of the best hit to a species outside of the ancestral lineage and *nbsA* is the normalized bit score of the best hit to a species within the ancestral lineage. AI scores range from -1 to 1, being greater than zero if the predicted protein sequence had a better hit to species outside of the ancestral lineage and can be suggestive of either HGT or contamination^33^. To identify HGTs present in multiple species, a recipient sub-lineage within the larger ancestral lineage may also be specified to identify their shared HGT candidates (Supplementary Fig. 3). All hits to the recipient lineage are skipped so as not to be included in the *nbsA* calculation. To identify candidate HGTs acquired from distant gene donors (e.g. viruses, bacteria, or plants) we first ran the AI pipeline using Ascomycota (NCBI:txid4890) and Xylariomycetidae (NCBI:txid 222545) as the ancestral and recipient lineages, respectively (Supplementary Fig. 3). To identify candidate horizontal transfers of genes predicted by antiSMASH to be in a SMGC from more closely related donors (e.g., other filamentous fungi), we ran the AI pipeline a second time using Xylariales (NCBI:txid 37989) as the ancestral lineage and manually curated subclades (see Supplementary Table 1) as recipient lineages (see Supplementary Fig. 3).

All HGT candidates were selected for tree building if they passed the following filters: (i) AI score of greater than 0, (ii) significant hits to at least 25 sequences in the custom database, and (iii) at least 50% of top hits were to sequences outside of the ancestral lineage. Full-length proteins corresponding to the top < 200 hits (E-value < 1 × 10^-3^) to each AI candidate were extracted from the custom database using esl-sfetch^108^. Sequences were aligned using MAFFT v7.407 using --auto to select the best alignment strategy^97^. The number of well aligned columns was determined with trimAL v.1.4. rev15 using its gappyout strategy^109^ only alignments with ≥ 50 retained columns after trimAL were retained for phylogenetic analysis. Phylogenetic trees were constructed using the untrimmed MAFFT alignment as input using IQ-TREE v1.6.10^98^ using the built in ModelFinder to determine the best-fit substitution model^110^ and performing SH-aLRT and ultrafast bootstrapping analyses with 1,000 replicates each. Phylogenies were visualized using iTOL v4^111^.

Although the case for HGT is strongest when genes suspected of being horizontally acquired have well-supported phylogenetic associations that contradict accepted species relationships^112^, our initial query-based trees often lacked sufficient taxon sampling to be incongruent with the accepted species phylogeny. This was frequently the case when evaluating candidate HGTs from distant donors. For example, a query-based gene tree may contain only the recipient Xylariales and bacterial sequences but lacked sequences from other filamentous fungi. Sequences from other fungi add important context to indicate where the Xylariales sequences would have grouped if inherited vertically. Therefore, for identifying high confidence transfers from distant donors (i.e., first AI analysis), we combined the AI and OrthoFinder analyses to construct trees containing homologous sequences from additional fungi. For orthogroups with one or more AI candidates, we combined all orthogroup sequences with all extracted top hits to each AI candidate. Sequences were aligned and trees constructed as described above. Each phylogenetic tree was then manually curated to verify HGT with either high or low confidence. To be considered a high confidence candidate, HGT events had to meet the following criteria: (i) the association between donor and recipient clades was supported by ultrafast bootstrap >= 95 and (ii) recipient clade consisted of sequences from two or more species. If the candidate met one of the two criteria, the HGT was considered low confidence. Phylogenies that that not meet these criteria were excluded.

### Statistical analyses

To assess whether genes within different functional categories are associated with ecological mode (endophytic and non-endophytic), we performed phylogenetically independent contrasts (PICs^113^) with the function ‘brunch’ of the package ‘caper’ version 1.0.1^114^ in R version 3.6.1 (Supplementary Table 6). All other statistics were done in R version 3.6.1 or JMP version 15.1 (SAS Institute Inc., Cary, NC).

### Data availability

Raw sequence data, assembled sequences, and genome annotations are available through the corresponding MycoCosm portal (https://mycocosm.jgi.doe.gov/). NCBI accession numbers for raw reads and assemblies are listed in Supplementary Table 1. All other data can be found in FigShare Repository (DOI 10.6084/m9.figshare.c.5314025).

## ACKNOWLEDGEMENTS

Funding for the project was provided by the DOE JGI Large-scale Community Science Project (Grant number 503506). MEEF was funded by the Office for Research, Innovation and Impact at the University of Arizona and the University of Arizona BIO5 Postdoctoral Fellowship Program. FL and JM acknowledge the financial support from the NSF DEB-1541548 and DEB-1046065. We thank F. Martin, P. Gladieux, J. Spatafora, R. Vilgalys, and K. ÒDonnell for permission to use unpublished JGI F1000 genomes; D. Bellomo, Y. Sanchez-Rosario, and S. Valdez for laboratory assistance; and the Genomics Analysis and Sequencing Core (GATC), the Arizona Genomics Institute (AGI), and the High-Performance Computer (HPC) at the University of Arizona for technical support.

## AUTHOR CONTRIBUTIONS

Designed research: JMU, JHW, AEA, MEEF; Performed field or laboratory research: JMU, LPM, YMJ, AEA, FL, JM; Contributed fungal isolates or analytic tools: YMJ, DCE, RM, JHW, JCS, KYC, JGI authors; Analyzed data: MEEF, JMU, JHW, KEE, KS, ZK, JGI authors; Wrote the paper: MEEF, JMU, JHW, with contributions from all authors.

## COMPETING INTERESTS

The authors declare no competing interests.

## SUPPLEMENTARY MATERIALS

### Functional annotation of orthogroups

All 1,451,488 genes from the 121 genomes (ingroup and outgroup) were clustered into 104,604 orthologous groups (i.e., orthogroups), and ca. 25% (26,825) were assigned functional annotations (Supplementary Table 7). Gene ontology (GO) terms were designated for 6,458 orthogroups (6.2%), while 10,820 (10.3%) and 11,144 (10.7%) of gene families were assigned to InterPro and Pfam domains, respectively. A small fraction of orthogroups were assigned IDs as carbohydrate-active enzymes (CAZyme; 720, 0.7%), peptidases and peptidase inhibitors (MEROPS DB; 443 orthogroups, 0.4%), and transporters (TCDB; 1154, 1.10%). The total number of gene families with signal peptides was 15,076 (14.4%), among which 2,869 (2.7%) were annotated as effectors (Supplementary Table 7). When all ingroup taxa were compared, we observed no significant differences between the number of genes in different functional annotation categories (e.g., CAZymes, transporters, etc.) and ecological mode (i.e., endophytic or non-endophytic; see Supplementary Table 6).

### Evolutionary relationships of endophytic, saprotrophic, and pathogenic Xylariaceae s.l. and Hypoxylaceae

Maximum likelihood phylogenomic analyses were performed with IQ-Tree using a concatenated matrix of 1,526 universal, single-copy orthogroups (Fig. 1a; Supplementary Fig. 1). Phylogenomic results support the monophyly of the newly proposed families of Graphostromataceae and Hypoxylaceae^23^, as well as previously observed relationships among genera^7^ (Supplementary Fig. 1). Dense gene sampling resulted in improved resolution and statistical support for deeper internal branches compared to a previous five-gene analysis^7^. Inclusion of previously unstudied endophytic taxa markedly increased the known phylogenetic diversity of the family^7^ (Supplementary Fig. 1), highlighting the importance of including unnamed endophytes (which are typically sterile mycelium in culture which precludes morphological characterization and formal naming; but see^115^) in phylogenetic studies.

Our analyses revealed seven endophytic isolates in five distinct clades (i.e., clades E2, E4, E5, E6, and E6) nested between the Graphostromataceae and Xylariaceae *sensu stricto*. To better ascertain their taxonomic relationships, we performed additional phylogenetic analysis that included recently published xylarialean taxa closely related to Xylariaceae and Graphostromataceae (i.e., *Barrmaelia*, Barrmaeliaceae^22^; *Linosporopsis* and *Clypeosphaeria*, Xylariaceae^116^). Briefly, we queried sequences of RPB2, alpha-actin, beta-tubulin, and ITS nrDNA for 35 taxa not included in previous multilocus analyses that contained xylarialean endophytes (ref ^7^) (e.g., *Barrmaelia*, *Linosporopsis, Clypeosphaeria Entosordaria, Graphostroma; Cryptostroma*^117^) against the reference multilocus Xylariaceae tree ^7^ in Tree-Based Alignment Selector Toolkit (T-BAS) v2.2 ^118–120^ with the evolutionary placement algorithm in RAxML^121^. The settings that we used to place taxa within the reference tree were as follows: UNITE filter off, no clustering, likelihood weights (fast), with the outgroup selected, and data were retained for all isolates. This analysis revealed that endophytes in clade E4 are sister to *Barrmaelia*, endophytes in clade E5 are sister to *Linosporopsis*, and endophytes in clade E6 are sister to *Clypeosphaeria* (Supplementary Fig. 1). Thus, our use of Xylariaceae *sensu stricto* and Xylariaceae *sensu lato* corresponds to ref ^22^ (see Supplementary Fig. 1).

### Phylogenomic results are robust to outgroup taxa, gene selection, and model of evolution

To assess the robustness of our phylogenetic results we reconstructed the phylogeny of Xylariaceae s.l. and Hypoxylaceae using four different approaches that differed in either outgroup taxon selection, model of inference, or orthologous gene set. First, we performed a maximum likelihood (ML) analysis of 1,526 single-copy orthogroups (found across all 121 ingroup and outgroup taxa) with the LG model of evolution (Fig. 1, Supplementary Fig. 1). Second, we performed an ML analysis of the same orthologous genes and 121 taxa, but with the JTT+F+I+G4 model of evolution, which was the best evolutionary model selected by ModelFinder in IQ-TREE (Supplementary Fig. 2). Third, we performed an ML analysis with the JTT+F+I+G4 model of evolution and the same orthologous genes, but after removing non-Xylariales taxa from the outgroup (data not shown). We performed a fourth ML analysis with all taxa (i.e., 121 ingroup and outgroup), but with 1,086 protein sequences identified as universal fungal orthologs with fungal genomes from JGI Mycocosm^62^. JGI orthologs were identified in genomes using the PHYling pipeline (DOI: 10.5281/zenodo.1257002; https://github.com/stajichlab/PHYling_unified). All phylogenetic analyses were performed with IQ-TREE multicore v1.6.1178 with 1,000 ultrafast bootstrap replicates (data not shown). All phylogenetic analyses resulted in similar topologies. However, relationships among taxa in the Xylaria HY and E9 clades differed slightly with the LG model (analysis 1) and the JTT+F+I+G4 (analysis 2) (see Supplementary Fig. 2).

In previous multi-locus analyses, the endophytic isolate Xylariaceae sp. FL2044 was placed as a sister to the Xylariaceae s.l. and Hypoxylaceae^7^. However, our concatenated phylogenomic analyses using different sets of single copy orthologs consistently placed FL2044 as basal within the monophyletic clade containing Xylariaceae s.l. (Fig. 1a; Supplementary Fig. 1). To confirm the placement of FL2044, we computed single-gene trees with IQ-TREE and used ETE Toolkit (http://etetoolkit.org/) to quantify the number of genes that supported the placement of FL2044 as recovered in our concatenated phylogenomic analyses. Overall, of the 882 single-gene trees where the placement of FL2044 was highly supported (i.e., >75% bootstrap), 297 (33.7%) agree with the placement of FL2044 in our concatenated analyses (Fig. 1; Supplementary Figs. 1,2). However, the placement of FL2044 in the Xylariaceae s.l. clade also was supported by network analysis of shared orthogroups (Supplementary Fig. 10).

### Determination of core, family-specific, clade-specific, and isolate-specific orthogroups and SMGCs

To visualize the distribution of orthogroups and SMGCs across taxa, we categorized orthogroups/SMGCs into 10 categories (see Supplementary Tables 3,7). To visualize the relative abundance of these categories across the phylogeny, we combined categories in the following manner for Supplementary Fig. 2e. Core: orthogroups/SMGCs present in all 121 taxa (cat a), as well as orthogroups/SMGCs present in all Xylariaceae s.l. and Hypoxylaceae taxa and in some outgroup taxa (cat c). Family-specific (i.e., Xylariaceae s.l. and Hypoxylaceae specific): orthogroups/SMGCs present in all or some Xylariaceae s.l. and Hypoxylaceae taxa, but absent in outgroup taxa (cat b and cat d). Hypoxylaceae-specific: orthogroups/SMGCs present in all or some Hypoxylaceae taxa, but absent in Xylariaceae s.l. taxa and outgroup taxa (cat e and cat f). Xylariaceae s.l.-specific: orthogroups/SMGCs present in all or some Xylariaceae s.l. taxa, but absent in Hypoxylaceae taxa and outgroup taxa (cat g and cat h). Isolate-specific: orthogroups/SMGCs found only in a single genome (cat j). Orthogroups/SMGCs that did not fall into any of these categories were defined as “other” (cat i). Examples of the “other” category include orthogroups/SMGCs that are present in some outgroup taxa, as well as some Hypoxylaceae and/or Xylariaceae s.l. taxa. Orthogroups/SMGCs distributions falling in the “other” category may have arisen through HGT, ancestral gene duplication and gene loss, or interspecific hybridization^122^. We found that no orthogroups were both unique to- and universally present in all Xylariaceae s.l. and Hypoxylaceae taxa (Supplementary Table 7d). A single orthogroup (annotated as a putative signaling peptide; OG0009755) was specific to and universally distributed in the Hypoxylaceae clade, but no orthogroups met these criteria for the Xylariaceae s.l. clade.

Overall, ca. 21-37% of the orthogroups per genome (mean = 27.4%) represented orthogroups shared by all 121 taxa (i.e., core genes; n =2,656 total) (Supplementary Fig. 2e; Supplementary Table 7). An additional 1,831 orthogroups were present in all Xylariaceae s.l. and Hypoxylaceae and one or more outgroup taxa (Supplementary Table 7d), representing an average of 14-23% orthologous gene families per genome (mean = 18.5%; Supplementary Fig. 2e). Gene families unique to Xylariaceae s.l. and Hypoxylaceae (i.e., absent in the outgroups and present in at least one genome in both Hypoxylaceae and Xylariaceae s.l. clades) represented, on average, ca. 1.6% of orthogroups per genome (Supplementary Fig. 2e, orange bars). An average of 3.0% and 3.8% of orthogroups were unique to Hypoxylaceae or Xylariaceae s.l. taxa, respectively (Supplementary Fig. 2e; Supplementary Table 7d).

Orthogroups unique to a single genome (i.e., dispensable orthogroups) represent ca. 1.4 to 15.6% of the orthogroups per genome for Xylariaceae s.l. and Hypoxylaceae (Supplementary Fig. 2e). Functional annotation using euKaryotic Orthologous Groups (KOGs) revealed a greater fraction of dispensable orthogroups were predicted to be involved in cellular processes and signaling (i.e., 42.6%) compared to core orthogroups (27.7%), including a higher fraction of orthogroups annotated as defense mechanisms and extracellular structures (Supplementary Fig. 13; Supplementary Table 7f). Dispensable orthogroups also were more likely than core orthogroups to encode proteins secreted through the general secretory pathway (15.0% vs 2.7%), supporting the hypothesis that strain-specific genes may provide ecological adaptations^44^. However, the functions of the majority of dispensable orthogroups remain unknown (i.e., only 20% had functional annotation vs. 90% of core orthogroups), similar to results from Dothideomycetes genomes^44^.

### Comparison of Hypoxylaceae and Xylariaceae s.l. SMGCs to MIBiG

Although there has been increasing biochemical characterization of metabolites from species of Xylariaceae s.l. and Hypoxylaceae (e.g., terpenes and polyketide compounds^10^), fewer studies have linked metabolites to gene clusters. Here, we compared predicted SMGCs to a reference database of known metabolites clusters (MIBiG^30^). Only 25% of predicted SMGCs (n = 1,711, belonging to 816 cluster families) had BLAST hits to 168 unique MIBiG^30^ accession numbers (Supplementary Table 3b). The majority of MIBiG hits were classified as PKSI (808 hits), terpene synthases (268 hits), and PKS-NRPS hybrids (253 hits). The remaining 382 hits were classified as NRPS, PKS-Other, RiPPS, and Other SMGCs. The average similarity of SMGCs to a MIBiG accession was 54% (range 13-100%) (Supplementary Table 3), but 587 xylarialean SMGCs were 100% similar to 38 MIBiG accessions (Supplementary Table 3).

Similarity to MIBiG is currently defined as the percentage of genes in an SMGC with significant BLAST hits to a known SMGC^39^, yet similarity can be difficult to assess given the dynamic nature of SMGCs (i.e., frequent gene duplications, gene losses, and HGT^41,123^) and the potential for *in silico* methods to misidentify cluster boundaries. For example, the griseofulvin cluster of *Penicillium aethiopicum* is predicted to contain 21 genes, but only core genes Gsf A, I, and G have been experimentally validated^40^. *Xylaria* taxa, despite lacking 13 genes (GsfR2, GsfK, GsfR1, GsfJ, GsfH and all eight genes of unknown function; Supplementary Fig. 4a), produce detectable levels of griseofulvin in culture (Supplementary Fig. 4b, see also ^124^). However, lower similarity may also reflect true differences in cluster composition and the production of similar, but distinct metabolites. Variation may also represent null alleles unable to synthesize the metabolite (e.g., aflatoxin in *A. flavus*^125^). Currently, databases such as MIBiG primarily contain metabolites from bioactive fungi with important roles as human or plant pathogens, and increased effort is needed to link metabolites from xylarialean fungi to specific gene clusters.

### Correlation between SMGC content and other functional categories

Consistent with the prevalence of SMGCs among clades of fungi known for their saprotrophic ecological roles (e.g., *Aspergillus*, *Penicillium*^13,14^), we observed that in genomes of non-endophytic Xylariaceae s.l. and Hypoxylaceae SMGC abundance is positively correlated with the number of genes important for saprotrophy (e.g., CAZymes, transporters) and putative pathogenicity (e.g., effectors, peptidases), even after accounting for differences among clades and genome sizes (P<0.01; Supplementary Table 6e). Our results are consistent with strong selection for saprotrophs to maintain large gene repertoires to degrade diverse lignocellulosic compounds^44^, as well as highly diverse SMGCs that likely increase competitive abilities in diverse microbial communities^11,49,50^. However, no such correlation was observed for genomes of endophytes in either clade, despite endophytes containing the same fraction of SMGC accessory genes annotated as CAZymes, peptidases, and effectors (but see Hypoxylaceae clade paired comparisons; Supplementary Table 6c).

### Intergenic distances, repetitive elements, effectors, and SMGCs

The software BEDTools version 2.29.2^126^ was used to calculate the distance between adjacent genes (intergenic distance) and the distance between each gene and the closest repetitive element on the 5’ and the 3’ end following ref ^127^. Results were visualized using the package ‘ggplot2’ version 3.3.2 in R and previously published code^127^ (https://github.com/lambros-f/blumeria_2017). The mean intergenic distance for all Xylariaceae s.l. and Hypoxylaceae genomes was 1,776 ± 415 bp. For all genomes, the distribution of intergenic distances followed a normal distribution, except for the genome of *Sodiomyces alkalinus,* which also displayed an increase in the frequency of genes with an intergenic distance towards 10,000 bp. Repetitive elements occurred more frequently in gene-sparse regions and at the end of contigs (Supplementary Table 2). Since *de novo* genome assemblers can collapse when reaching a repetitive region larger than the read length itself ^128^, we surmise that our genome assemblies may be fragmented because of complex regions rich in repetitive elements.

To identify whether SMGCs and effectors were in regions of the genome with high repeats and sparse gene content, we performed the same calculation of intergenic distances and visualized the locations as a function of gene density and TE location. We observed no significant differences in the density of repetitive elements for effector genes vs. non-effector genes for genomes of Hypoxylaceae or Xylariaceae s.l. taxa (Supplementary Fig. 5). In the majority of Xylariaceae s.l. and Hypoxylaceae genomes, numerous SMGCs, and genes annotated as effectors are located at the edge of contigs in gene sparse/high repeat regions including the griseofulvin cluster in *Xylaria*. sp. However, there was no relationship between SMGC number (residuals after accounting for genome size) and the number of scaffolds obtained from genome assembly (Supplementary Fig. 16) suggesting that fragmentation of genome assemblies did not artificially increase the predicted number of SMGCs^103^. Repetitive-rich regions, often near telomeres and centromeres, can represent hotspots of gene gain/loss events as transposable elements facilitate gene dispersal both within and among genomes^11,17,129^. The presence of SMGCs in these regions may drive the hyperdiversity of SMGCs within the Xylariales, as well as the discontinuous phylogenetic distribution of SMGCs across the studied genomes (see Figs. 1,3).

### Confirmation of griseofulvin HGT

We examined regions flanking in *Xylaria* sp. with and without the griseofulvin cluster to further confirm HGT. Briefly, 30 kbp sequences located up- and downstream of the griseofulvin cluster of *Xylaria flabelliformis* CBS 123580 were queried with BLASTn against closely related genomes without the griseofulvin cluster to identify homologous regions (*X. longipes* CBS 148.73 scaffold 57 and *X. acuta* CBS 122032 scaffold 139). Scaffolds containing these homologous regions, along with the scaffolds containing the griseofulvin cluster in *X. flabelliformis* NC1011 (scaffold 71), *X. flabelliformis* CBS 124033 (scaffold 75), *X. flabelliformis* CBS 123580 (scaffold 16), *X. flabelliformis* CBS 114988 (scaffold_56), *X. flabelliformis* CBS 116.85 (scaffold 29), were then aligned using Mauve^130^. In *X. flabelliformis* isolates, the scaffold alignment contains the up- and downstream homology blocks with the intervening griseofulvin cluster. Up- and downstream homology blocks were also found in *X. longipes* CBS 148.73; however, the griseofulvin cluster was not present, thus supporting the HGT of griseofulvin cluster in some taxa.

### Comparison of leaf litter decomposition among clades

To assess the ability of Xylariaceae s.l. and Hypoxylaceae fungi to degrade lignocellulose, we collected fresh, healthy, green leaf material from two individuals of *Quercus virginiana* and *Pinus halepensis* at the University of Arizona campus arboretum. Trees are cultivated in a park-like setting with supplemental water and appear healthy. For both species, leaves were washed in tap water to remove any surface debris. Washed leaves were autoclaved for 20 min to inactivate endogenous microbes and then dried overnight at 60°C. Autoclaved leaves (0.5 g) were placed into individual, sterile 100 mm Petri plates (three replicate plates per leaf substrate type for each fungal isolate). For each fungal isolate, a 6 mm plug of mycelium (actively growing on 2% MEA) was briefly homogenized with a sterile minipestle in 1 mL of sterile water until mycelia had visually separated from the agar chunks. From this 1 mL mixture, 75 µL was diluted with 3 mL of sterile water and mixed via pipetting to create the fungal inoculum. One mL of the diluted ground mycelium was placed directly on the sterile leaf surface in each Petri dish. Negative control samples were inoculated in parallel with sterile water. In total, we inoculated three replicate plates per fungal isolate per plant species (total of 120 plates). Petri plates were sealed with Parafilm and weighed on an analytical balance (mass_original_). Plates were stored in the dark at 26°C for the duration of the experiment (12 weeks). Each plate was weighed weekly, and the percent of leaf tissue covered with mycelium was visually scored (0 = no visible growth; 1 = 1-25% leaf coverage; 2 = 26-50% leaf coverage; 3 = 51-75% leaf coverage; 4 = 76-100% leaf coverage). Negative controls did not display fungal growth. We calculated mass loss for each replicate and control as mass_final_ = mass_week12_ –mass_original_. To account for water loss due to evaporation, we then subtracted the average value of the negative control plates (mass_norm_ = mass_final_ –mass_control_). We compared the normalized mass loss among clades with ANOVA (Supplementary Fig. 11).

### Metabolite extraction and identification

To induce the production of SMs and potentially verify SMGCs, we performed co-culture experiments with three isolates: *X. flabelliformis* NC1011, *Xylaria arbuscula* FL1030, and *Daldinia* sp. FL1419. Isolates were grown on *Aspergillus* defined media^131^. After one week, we removed 6 mm diameter plugs of actively growing mycelium from each isolate for three pairwise combinations of co-culture plates (i.e., NC1011 vs. FL1419; FL1419 vs. NC1030; NC1030 vs. NC1011). Briefly, agar plugs of two isolates were placed ∼4.5 cm apart across the horizontal diameter of a 100 mm Petri dish (see Supplementary Fig. 4b). We incolulated four replicate co-culture plates for each combination (total 12 interaction plates) and four plates containing each isolate alone (total 12 positive control plates). Plates were incubated at room temperature for 8-10 days or until the mycelium from the two isolates was ∼1cm apart. Using a sterile transfer tube, we harvested five 6 mm plugs of agar either (i) next to a single culture (i.e, positive control plates); (ii) in the space between isolates (i.e., interaction plates) to ensure the capture of exogenous SMs; or (iii) in the middle of media control plate. After harvesting agar plugs were placed into sterile, 2.0 mL microcentrifuge tubes, flash frozen in liquid Nitrogen, and stored at -80°C. Frozen samples were shipped to JGI for extraction and stored and -80°C until processed.

To extract metabolites for LC-MS/MS, samples were lyophilized dry (FreeZone 2.5 Plus, Labconco), then bead-beaten to a fine powder with a 3.2 mm stainless steel bead for 5 seconds (2x) in a bead-beater (Mini-Beadbeater-96, BioSpec Products). For extraction, 500 µL of MeOH was added to each sample, briefly vortexed, sonicated in a water bath for 5 minutes, and centrifuged for 5 min at 5000 rpm to pellet agar and cellular debris. The supernatant was transferred to a 2 mL Eppendorf, dried in a SpeedVac (SPD111V, Thermo Scientific), and stored at -80 °C. Extraction controls were prepared similarly but using empty tubes exposed to the same extraction procedures. In preparation for LC-MS/MS analysis, dried extracts were resuspended by adding 300 µL methanol containing 10 µg/mL of 2-Amino-3-bromo-5methylbenzoic acid (#R435902, Sigma) as internal standard, vortexed briefly, sonicated in a water bath for 10 min, and centrifuged (5 min at 5000 rpm). After centrifugation, 150 µL of the resuspended extract was filtered via centrifugation (2.5 min at 2500 rpm) through a 0.22 µm filter (UFC40GV0S, Millipore) and transferred to a glass autosampler vial.

Samples were analyzed on a system consisting of an Agilent 1290 UHPLC coupled to a Thermo QExactive Orbitrap HF (Thermo Scientific, San Jose, CA) mass spectrometer. Reverse phase chromatography was performed by injecting 2 µL extract into a C18 column (Agilent ZORBAX Eclipse Plus C18, 2.1×50 mm, 1.8 µm) warmed to 60°C with a flow rate of 0.4 mL/min equilibrated with 100% buffer A (100% LC-MS water with 0.1% formic acid) for 1 minute, followed by a linear gradient to 100% buffer B (100% acetonitrile w/ 0.1% formic acid) for 7 minutes, and then held at 100% B for 1.5 minutes. MS and MS/MS data were collected in both positive and negative ion mode, with full MS spectra acquired ranging from 90-1350 *m/z* at 60,000 resolution, and fragmentation data acquired using an average of stepped collision energies of 10, 20 and 40 eV at 17,500 resolution. Orbitrap instrument parameters included a sheath gas flow rate of 50 (au), an auxiliary gas flow rate of 20 (au), sweep gas flow rate of 2 (au), 3 kV spray voltage, and 400 °C capillary temperature. Sample injection order was randomized and an injection blank of methanol only run between each sample. Metabolites were identified based on comparing exact mass (ppm difference between detected m/z to a compound’s theoretical m/z) and comparing experimental MS/MS fragmentation spectra to that of standards. These data confirmed the production of griseofulvin by NC1011 when grown in co-culture with FL1419 (Supplementary Fig. 4b).

## Supplementary Tables

**Supplementary Table 1. (a)** Information for the 121 genomes included in this study; (b) Genome and assembly information for 96 Xylariaceae s.l. and Hypoxylaceae genomes included in this study.

**Supplementary Table 2.** RepeatMasker, RepeatScout, and RepBase Update classification of repetitive elements for 96 genomes of Xylariaceae s.l. and Hypoxylaceae.

**Supplementary Table 3.** (**a**) Secondary metabolite gene cluster (SMGC) annotations for the 121 genomes included in this study (according to antiSMASH) and grouped into families with BiG-SCAPE; (**b**) Distribution and percent similarity of Xylariaceae s.l. and Hypoxylacae SMGCs to 168 MIBiG accessions; (**c**) Count and percentage of all SMGCs and SMGC families per category (A-J); (**d-j**) Count and percentage of SMGCs per type (e.g., NRPS, Terpene, Other PKS, PKS-NRP Hybrids, Other, RiPP) per category (A-J).

**Supplementary Table 4.** (**a**) Count of catabolic gene clusters (CGCs) by anchor gene; (**b**) Presence/Absence of CGC families per genome; (**c**) Composition of the CGC families; (**d**) Genomic position and annotation of CGCs.

**Supplementary Table 5.** (**a**) Taxonomic and phylogenetic information for 4,262 putative HGT candidate genes identified by Alien Index (AI); (**b**) Manual curation of phylogenetic trees reveals 168 HGT candidates (each row is a unique transfer event; orthogroups may appear more than once); (**c**) Distribution of HGT counts per genome (HGT001-HGT-129 are high confidence transfers and HGT130-HGT290 are ambiguous transfers); (**d**) Functional annotation of 1,148 SMGC genes identified by the second Alien Index as candidate HGTs.

**Supplementary Table 6. (a)** Number of genes annotated as MEROPS, CAZymes, PCWDCs, SMGCs, CGCs, and putative HGTs for genomes of 96 Xylariaceae s.l. and Hypoxylaceae; (**b**) Statistical comparison between Xylariaceae s.l. and Hypoxylaceae genomes; (**c**) Statistical comparison between endophytic and non-endophytic genomes with phylogenetic independent contrasts (PICS); (**d**) Statistical analysis of genomic features for paired endophyte/non-endophyte sister taxa using least-squares means contrasts; (**e**) Pearson correlation of genomic features as a function of ecological mode and clade.

**Supplementary Table 7. (a)** Orthogroup summary statistics; (**b**) Orthogroup annotations; (**c**) Count and percentage of orthogroups and proteins per orthogroup category (A-J). (**d**) Orthogroups that comprise each category (A-J).

**Additional Files (**Available on FigShare Repository; DOI 10.6084/m9.figshare.c.5314025)

**Additional file 1.** InterProScan annotations for 96 Xylariaceae s.l. and Hypoxylaceae genomes.

**Additional file 2.** AntiSMASH output for the 96 Hypoxylaceae and Xylariaceae s.l genomes.

**Additional file 3.** Tables summarizing the ancestral gene reconstruction by Count v10.04. The ancestral gene content was reconstructed for the entire data set, as well as for subsets of orthologous gene families corresponding to different functional groups including (i) CAZymes; (ii) plant cell wall degrading CAZymes (PCWDCs); (iii) PCWDCs involved in the degradation of cellulose, hemicellulose, lignin, pectin, starch and inulin; (iv) peptidases; (v) peptidase inhibitors; (vi) transporters; (vii) transporters involved in the exchange of carbohydrates; (viii) transporters involved in the exchange of amino acids; (ix) transporters involved in the exchange of lipids; (x) transporters involved in the exchange of nitrogen; and (xi) effectors.

**Additional file 4.** Graphs of intergenic distances for each genome of Xylariaceae s.l. and Hypoxylaceae, overlaid with the location of secondary metabolite gene clusters, repeat elements, and effector genes.

**Additional file 5.** Graphs depicting the frequency of repetitive elements surrounding genes for each genome of Xylariaceae s.l. and Hypoxylaceae.

**Additional file 6.** Phylogenomic trees inferred by maximum-likelihood under the JTT+F+I+G4 model for (a) the whole dataset of 121 taxa and 1,526 protein sequences; (b) a subset of Xylariales taxa only and 1,526 protein sequences; and (c) the entire dataset of 121 taxa and 1,086 protein sequences.

**Additional file 7.** Table showing the sister clades to Xylariaceae sp. FL2044 recovered by the phylogenetic analysis of each of the 1,526 single-copy orthologous genes.

**Additional file 8.** Alignment of regions flanking the griseofulvin cluster in *Xylaria* sp. (a) Mauve^130^ alignment of the scaffolds containing the griseofulvin cluster in *X. flabelliformis* NC1011, *X. flabelliformis* CBS 124033, *X. flabelliformis* CBS 123580, *X. flabelliformis* CBS 114988, *X. flabelliformis* CBS 116.85, and scaffolds of the closely related *Xylaria longipes* CBS 148.73 and *Xylaria acuta* CBS 122032 showing similarity to the griseofulvin flanking regions of *X. flabelliformis* CBS 123580. (b) Same alignment after hiding the scaffolds of *Xylaria acuta* CBS 122032, *X. flabelliformis* CBS 124033, *X. flabelliformis* NC1011, *X. flabelliformis* CBS 114988. Locally collinear blocks are shown in the same colors. The plot inside the blocks indicates the level of sequence similarity. The ruler above each scaffold represents the nucleotide positions. The white boxes below represent coding sequences. The griseofulvin cluster is highlighted in light blue for *X. flabelliformis* CBS 123580; the purple block contains the griseofulvin protocluster.

---

**Supplementary Figure 1.**
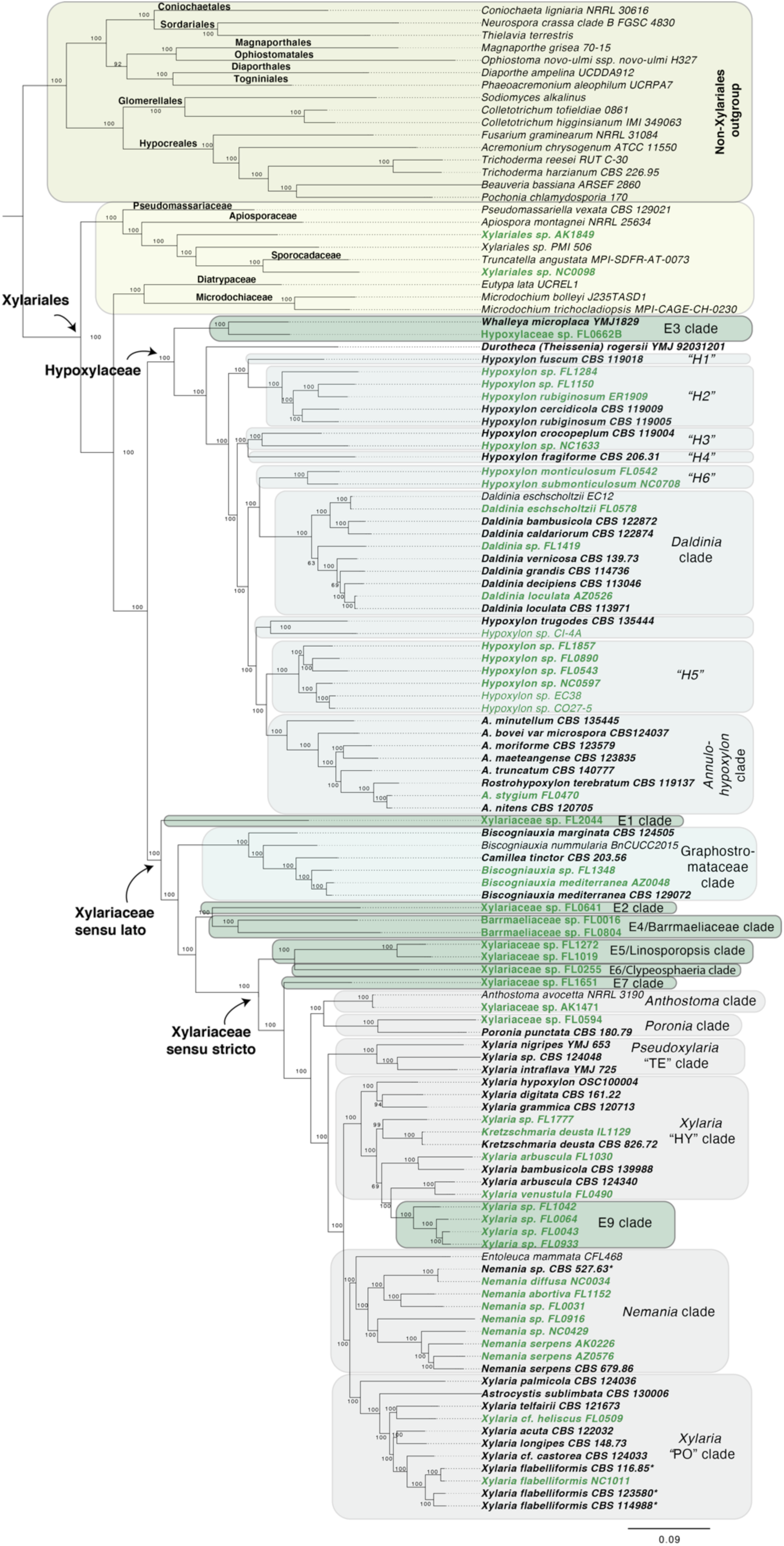
Phylogenomic tree inferred by maximum likelihood based on a combination of 1,526 universal single-copy orthologous protein sequences. Twenty-five Sordariomycetes species outside Xylariales were used as the outgroup (Supplementary Table 1a). Isolates sequenced in this study are highlighted in bold. Endophytes (i.e., fungi isolated from living, photosynthetic tissues of plants and lichens^7^) are indicated in green. Clade information is based on previously published studies (see refs ^7,22,23,42,43^). Numbers at nodes indicate ultrafast bootstrap support values from IQ-TREE^98^. The scale bar corresponds to the number of substitutions per site.

**Supplementary Figure 2.**
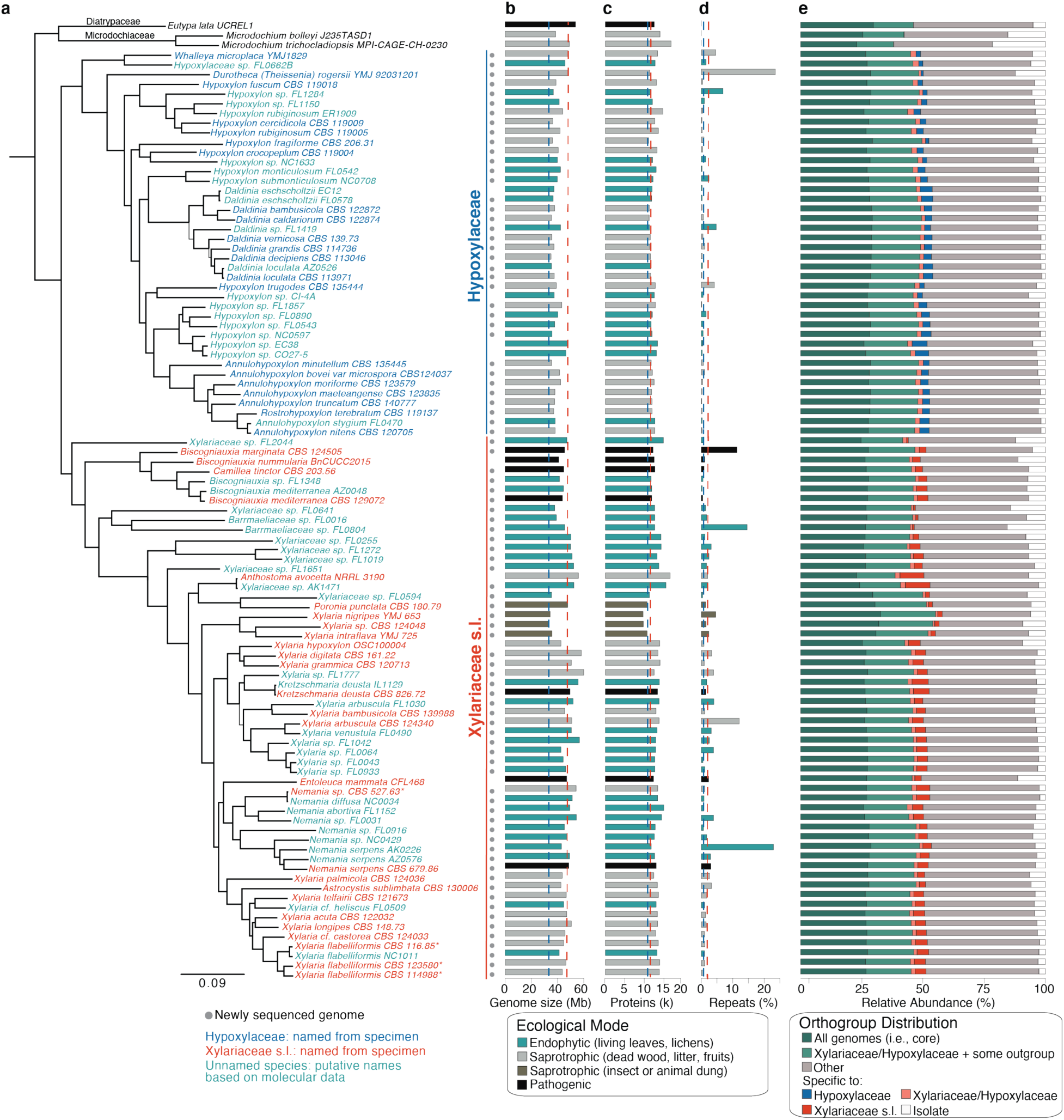
Phylogenomic reconstruction of Xylariaceae s.l. and Hypoxylaceae and genome statistics. (**a**) The maximum likelihood phylogram is based on 1,526 single-copy orthologous genes present in all genomes. Bootstrap values are shown in Supplementary Fig. 1. The scale bar indicates the number of substitutions per site. Names of reference taxa are colored according to their clade affiliation (dark blue: Hypoxylaceae; red: Xylariaceae s.l.). Undescribed endophyte species, putatively named based on phylogenetic analyses^7^, are shown in teal blue; (**b**) genome size; (**c**) predicted protein coding genes; and (**d**) percent transposable element (TE) content (bar colors correspond to ecological mode; see legend). Averages per major clade are shown with dotted lines in panels a-d; (**e**) relative abundance of core, family-specific, clade-specific, and isolate-specific orthogroups (see legend; Supplementary Table 3d).

**Supplementary Figure 3.**
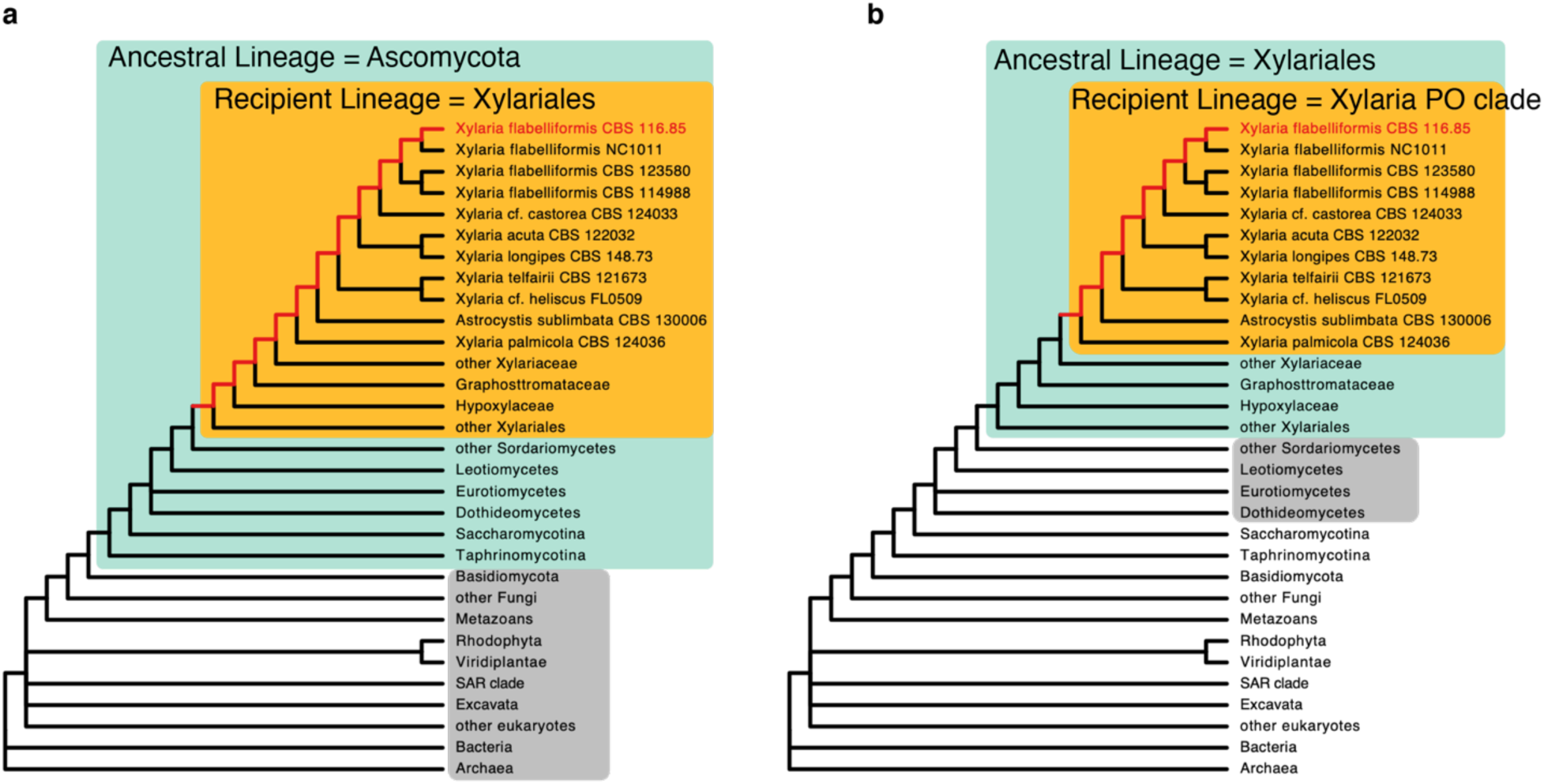
Overview of Alien Index (AI) calculations to identify HGT. In this example, *Xylaria flabelliformis* CBS 116.85 is the query genome. (**a**) AI screen to identify HGT candidates from more distant gene donors (grey box); candidates must have a better hit to sequences outside the ancestral lineage (Ascomycota; green box). By skipping all sequences to other Xylariales (orange box), HGT candidates could have been acquired at any point back to their last common ancestor (red branches) (**b**) AI screen to identify more recently acquired HGT candidates from other filamentous fungi (grey box). For this screen, candidates must have a better hit to sequences outside the Xylariales (green box). All sequences to other *Xylaria* “PO” clade were skipped (orange box) to identify shared HGT candidates acquired at any point back to the last common ancestor of the clade (red branches).

**Supplementary Figure 4.**
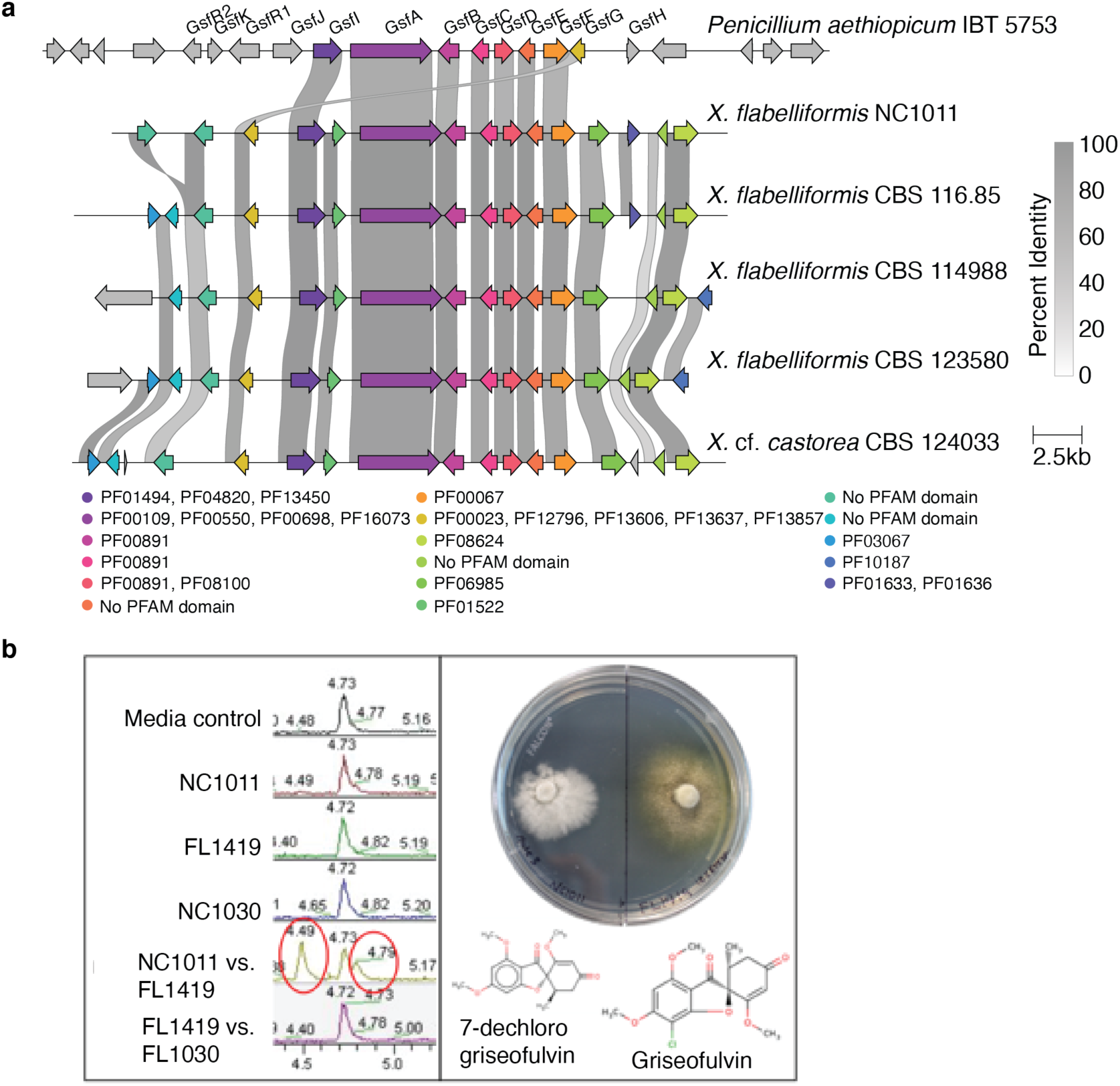
Similarity of the griseofulvin SMGC in *Penicillium* and *Xylaria* supports HGT. **(a)** Comparison of the griseofulvin cluster from *Penicillium aethiopicum* IBT 5753 (top) to five newly sequenced *Xylaria* genomes. Homologous genes are colored by PFAM domain. Connecting ribbons indicate percent amino acid identity to genes in the *Penicillium* cluster; (**b**) Metabolomic analysis of pairwise comparisons of *X. flabelliformis* NC1011, *Xylaria arbuscula* FL1030, and *Daldinia* sp. FL1419 illustrates production of griseofulvin by NC1011 during the interaction with FL1419, but not when grown alone or with isolate FL1030.

**Supplementary Figure 5.**
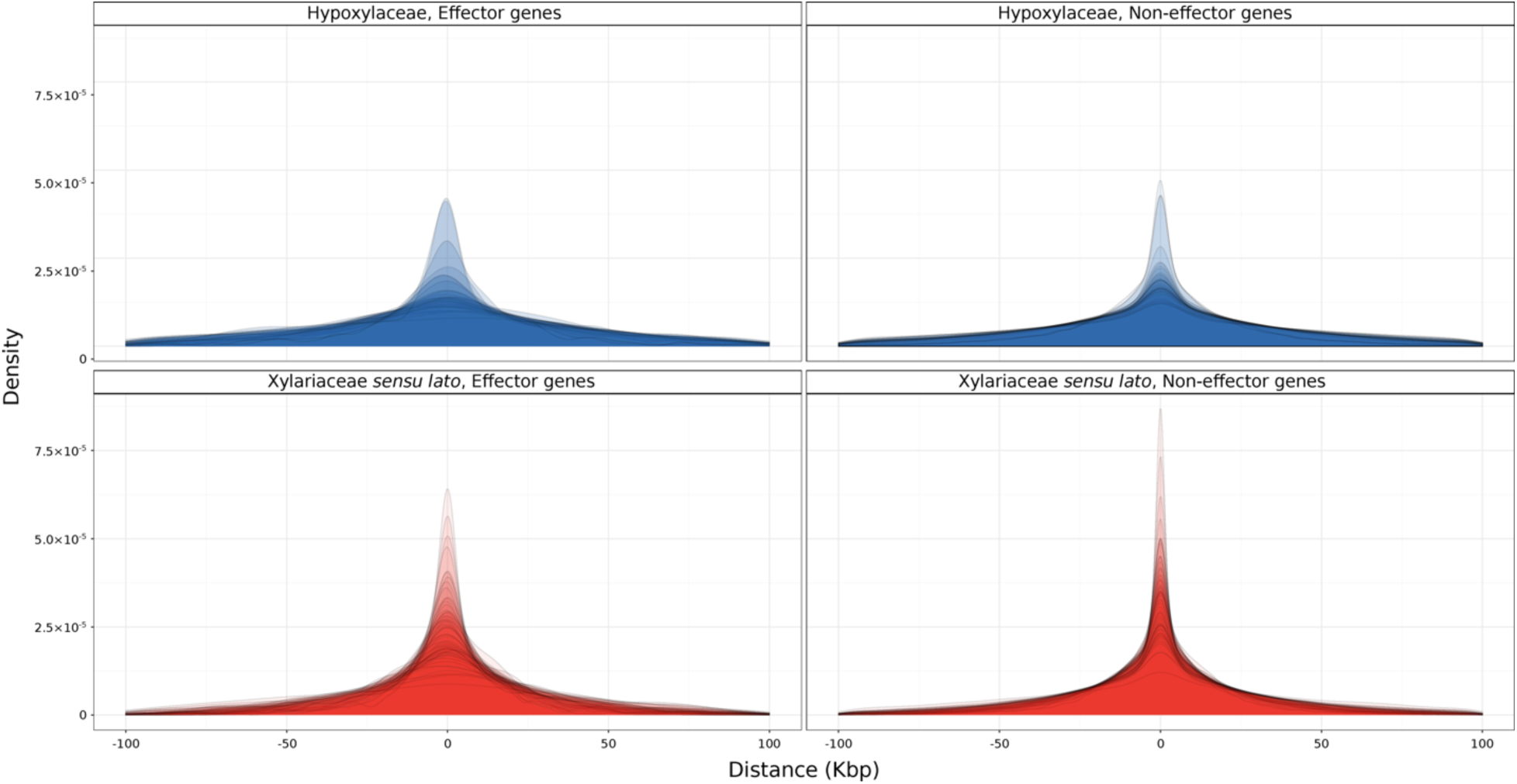
The density of repetitive elements surrounding genes was higher for Xylariaceae s.l. than for Hypoxylaceae genomes. Overlapped density plot of taxa per clade, illustrating the distance of repetitive elements from genes annotated as Effector (left) vs. Non-Effector genes (right). Negative distances indicate repetitive elements are located upstream of genes, while positive distances indicate repetitive elements downstream of the genes. Repetitive elements were identified by RepeatScout and RepeatMasker. Effector genes were predicted by EffectorP 2.0. The distances were computed using BEDTools v2.29.2.

**Supplementary Figure 6.**
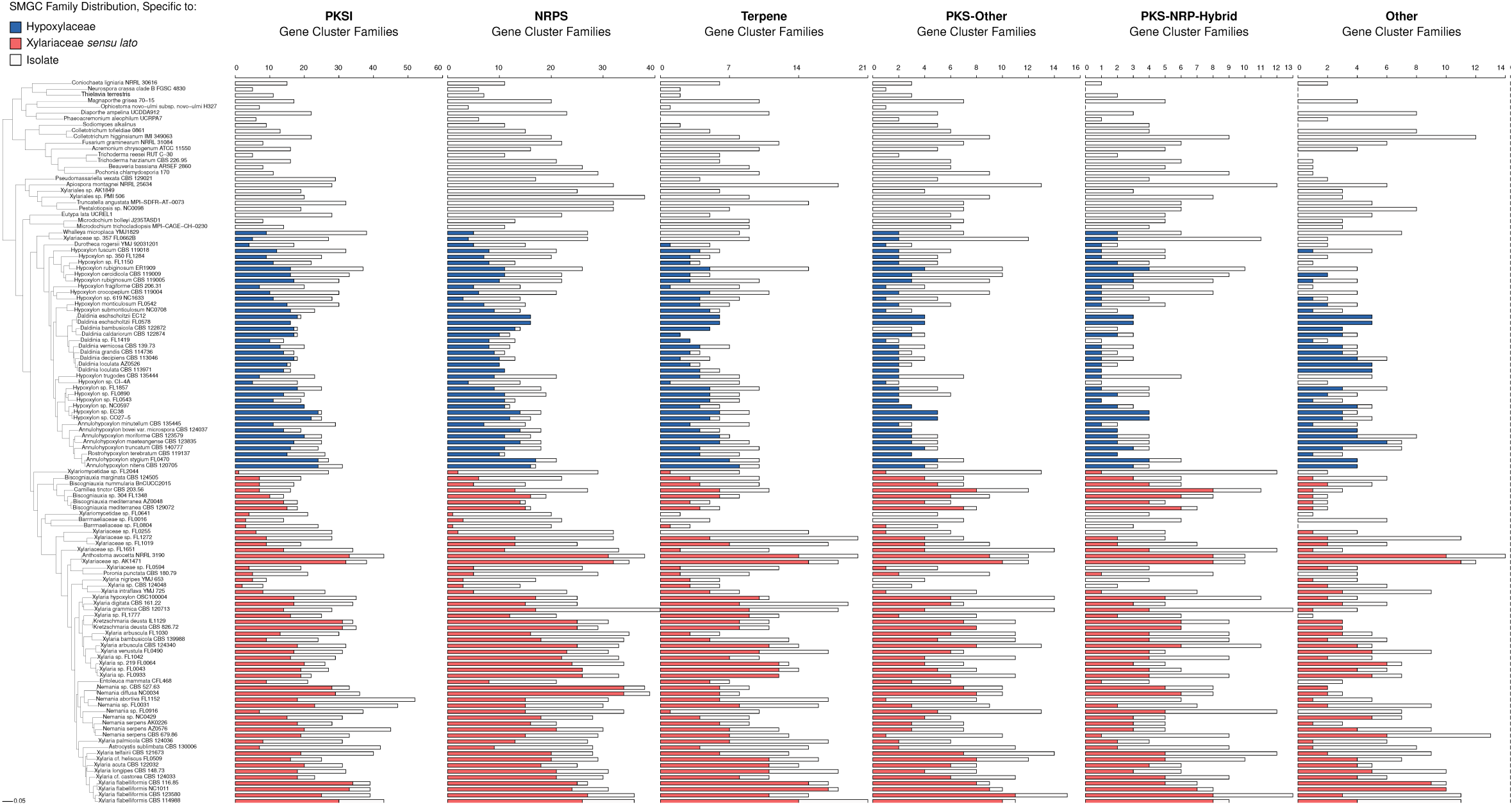
The majority of SMGCs are specific to Hypoxylaceae or Xylariaceae s.l. clades or individual isolates regardless of SMGC type. Phylogenomic tree of Xylariaceae s.l. and Hypoxylaceae and outgroup taxa with bar plots illustrating the number of SMGC families per genome, as well as the percentage of clade-specific and isolate-specific SMGC families for (**a**) PKSI; (**b**) NRPS; (**c**) Terpene; (**d**) PKS-Other; (**e**) PKS-NRP Hybrid; and (**f**) Other.

**Supplementary Figure 7.**
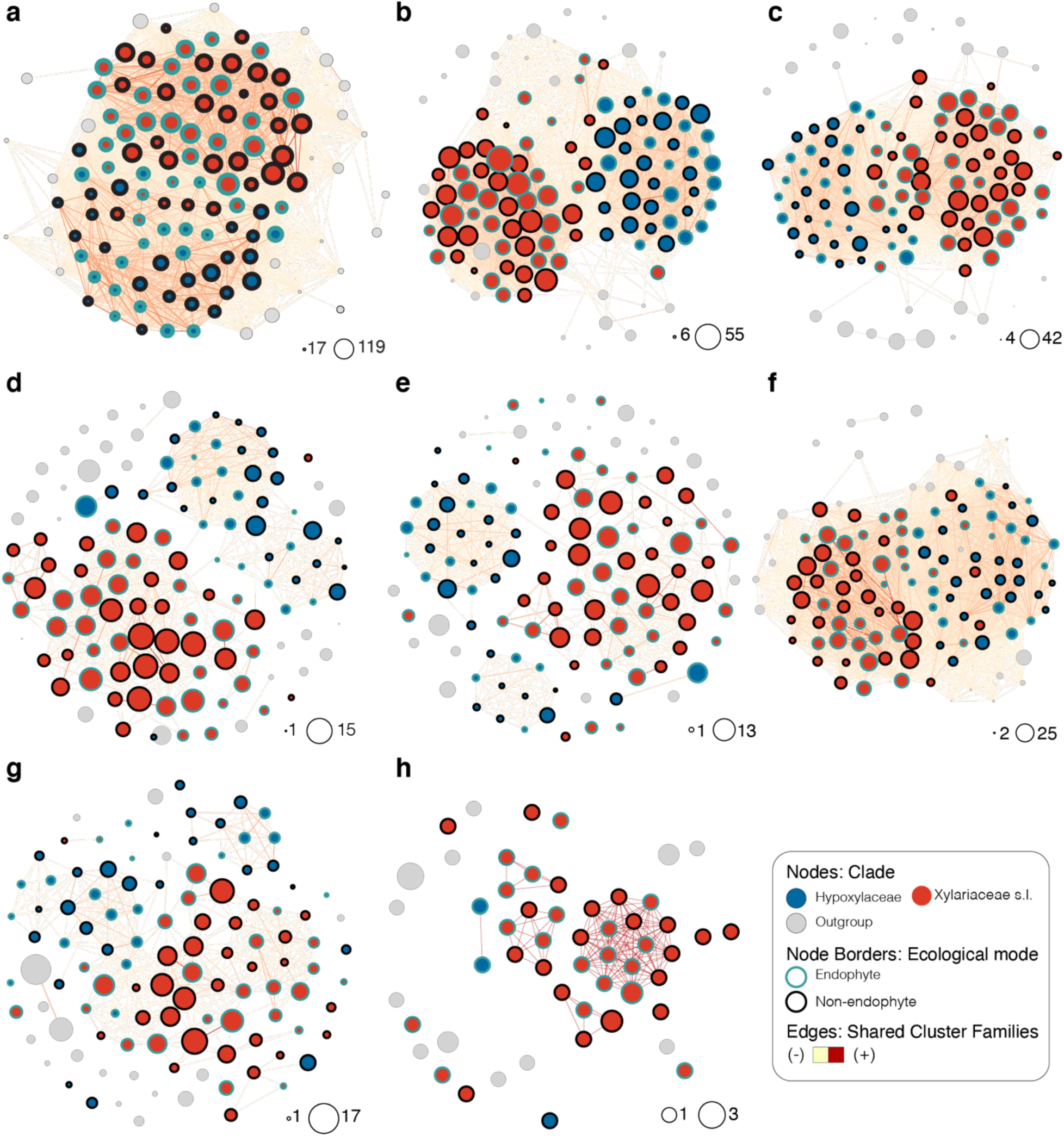
Network analysis illustrates the importance of clade rather than ecological mode for SMGC content. Network representation of SMGCs clustering from BiG-SCAPE. Each node represents the SMGC content per genome for (**a**) all SMGCs and SMGC sub-types: (**b**) PKSI; (**c**) NRPS; (d) PKS other; (e) PKS-NRPS Hybrids; (f) terpenes; (g) other; and (h) RiPPs. Networks are scaled by the count of gene clusters and positioned by a force-directed layout algorithm. Edges between two nodes are weighted by the number of shared clusters. Node color corresponds to clade. Nodes representing endophytic isolates are shown with blue borders.

**Supplementary Figure 8.**
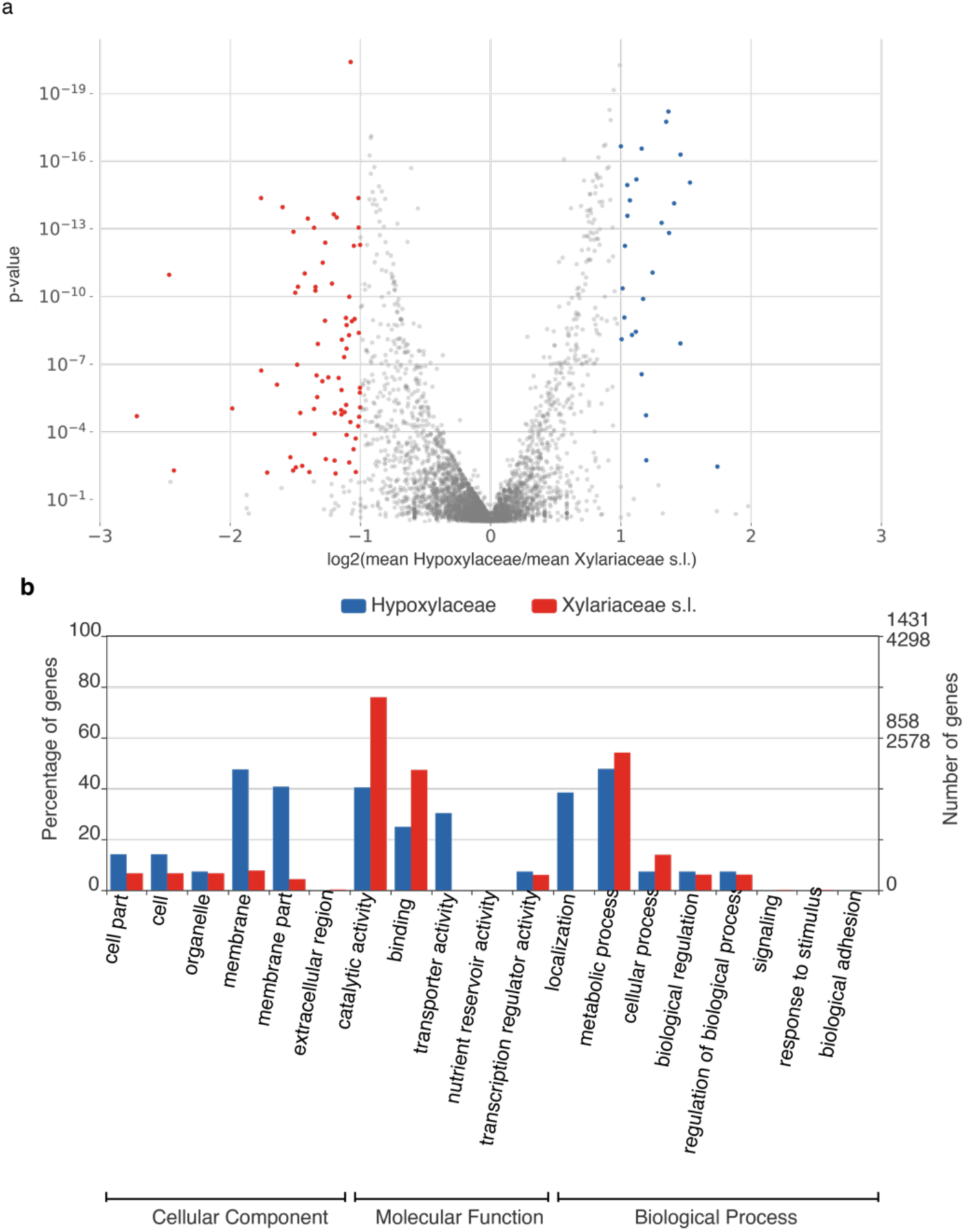
Twenty-six orthogroups were significantly enriched in the Hypoxylaceae clade, while 74 orthogroups were significantly expanded in the Xylariaceae s.l. clade. (**a**) Volcano plot of the protein count representation tests for orthogroups shared between the Hypoxylaceae and Xylariaceae s.l. clades. Orthogroups significantly enriched in Xylariaceae s.l. taxa are colored in red, while orthogroups significantly enriched in Hypoxylaceae taxa are colored in blue. Two-sided Mann-Whitney U-tests, p-value ≤ 0.01 and |log2FC| ≥ 1. (**b**) Comparison of enriched GO terms (level 2) of orthogroups significantly enriched in Hypoxylaceae taxa (blue) vs. Xylariaceae s.l. taxa (red). GO terms were analyzed and visualized using Web Gene Ontology Annotation Plot 2.0 (WEGO). See also Supplementary Table 3f for KOG annotation of enriched orthologs. The two-sided Mann-Whitney U-test was performed using SciPy^132^ through KinFin v1.0^99^.

**Supplementary Figure 9.**
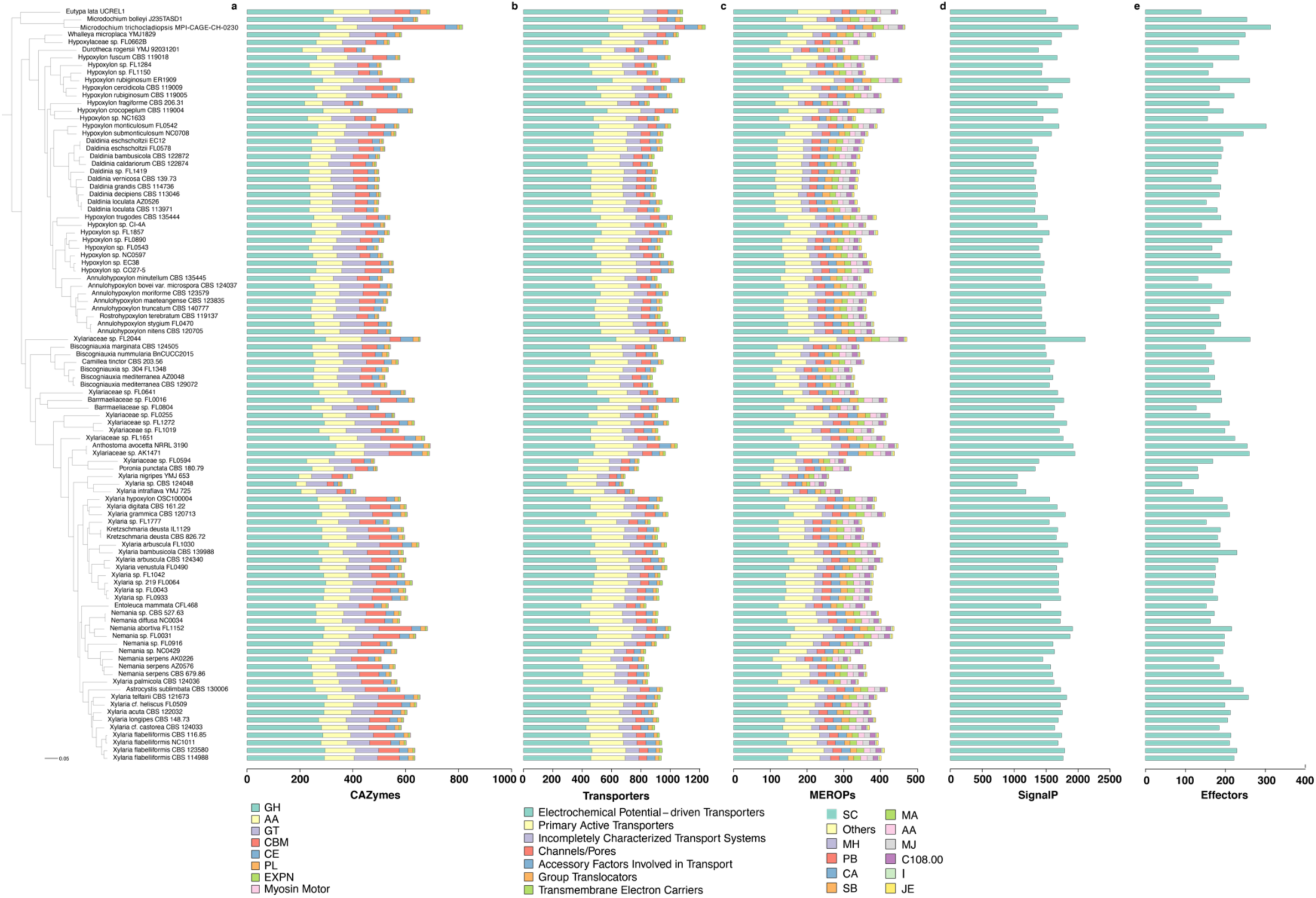
Relative abundance of functional gene categories across Xylariaceae s.l. and Hypoxylaceae. Phylogenomic tree and bar plot showing the abundance and identity of (**a**) carbohydrate-active enzymes (CAZyme); (**b**) peptidases and their inhibitors (MEROPs); (**c**) transporters (TCDB); (**d**) secreted proteins (SignalP); and (**e**) effectors (EffectorP). Colors refer to different classifications within each database (see legends).

**Supplementary Figure 10.**
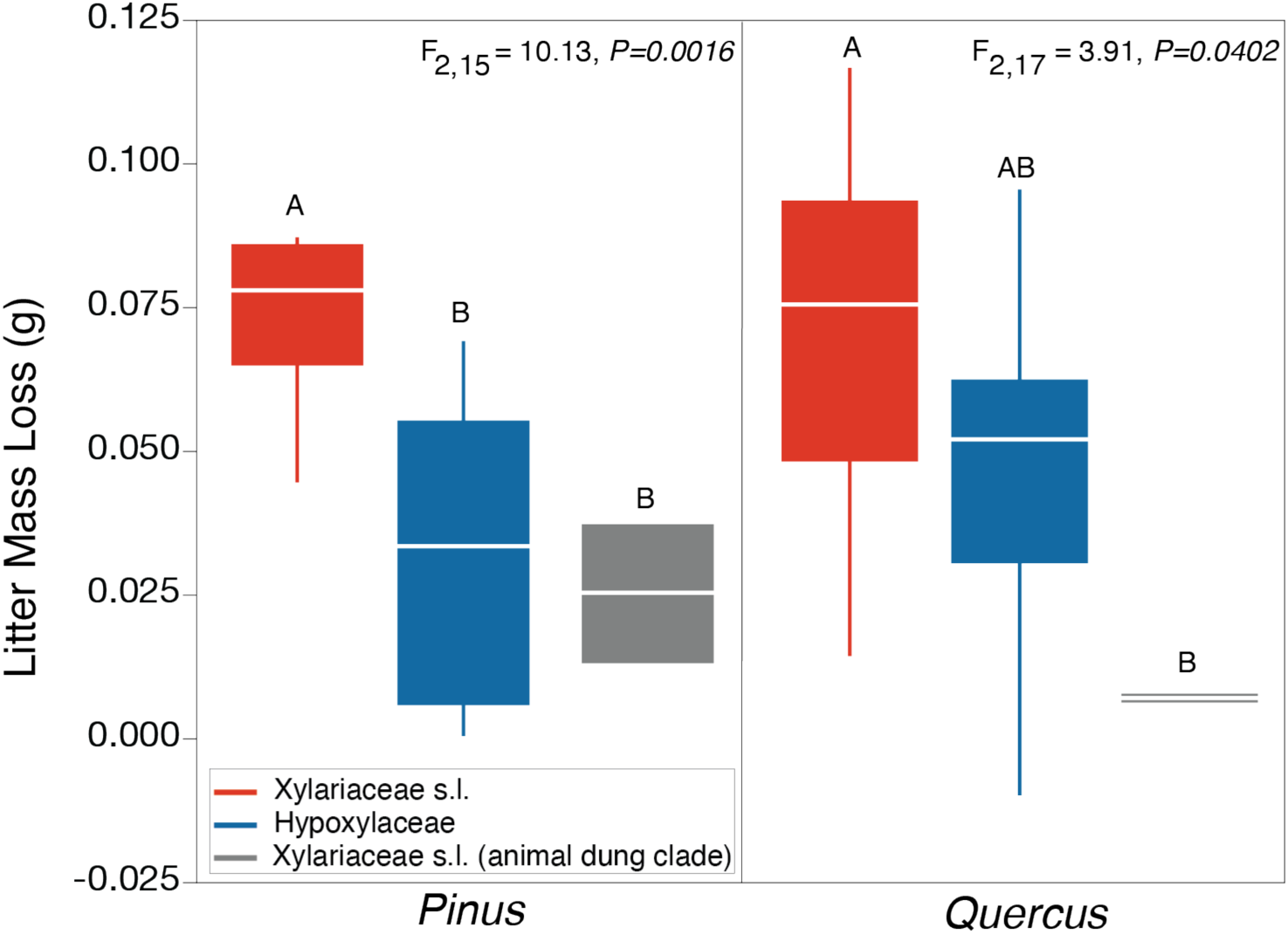
Xylariaceae s.l. taxa demonstrate increased decomposition abilities (estimated via mass loss) on leaf litter compared to fungi with reduced genomes (i.e., Hypoxylaceae and animal dung Xylariaceae s.l. in the *Poronia* clade). Interquartile box plots showing median and interquartile range. We observed significant differences among means of each clade on both *Pinus* and *Quercus* leaves (ANOVA). Letters indicate significant differences after post-hoc Tukey’s HSD. See Supplementary Table 1 for a list of isolates included in the mass loss experiment.

**Supplementary Figure 11.**
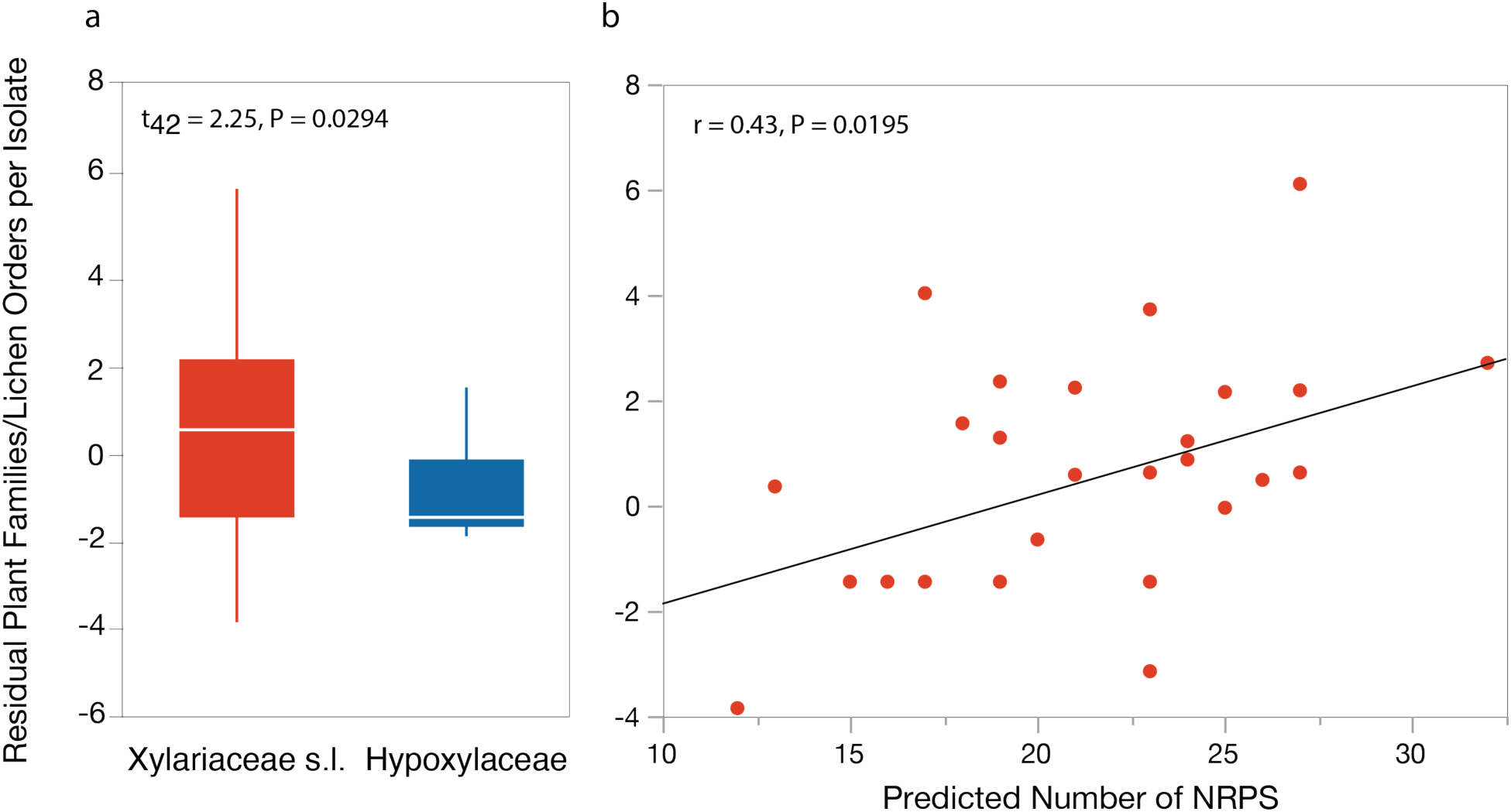
Endophytes in the Xylariaceae s.l. clade have greater host breadth, which is correlated with an increase in NRPS SMGCs. **(a)** A quantile box plot showing the interquartile range and median of endophyte host breadth (measured as total number of plant families and lichen orders with which a fungal OTU was cultured; see ^7^) as a function of major clade (color). T-test comparison of means illustrates greater host associations for Xylariaceae s.l. taxa (t-test, t_42_ = 2.25, P= 0.0294). A similar pattern was observed when only the number of plant families are compared (Wilcoxon: *χ*^2^ = 4.14, P=0.0413), but not lichen orders (Wilcoxon: *χ*^2^ = 1.77, P=0.1834). **(b)** Relationship of Xylariaceae s.l. endophyte host breadth and the number of SMGCs classified as NRPS. A similar pattern was observed when only the number of lichen orders was used to estimate host breadth (Pearson correlation: r = 0.4944, P = 0.0064), but not for the number of plant families. Host breadth is also correlated with the number of HGT events (r = 0.43, P = 0.0193), but HGT events and SMGC content are not independent (Fig. 6).

**Supplementary Figure 12.**
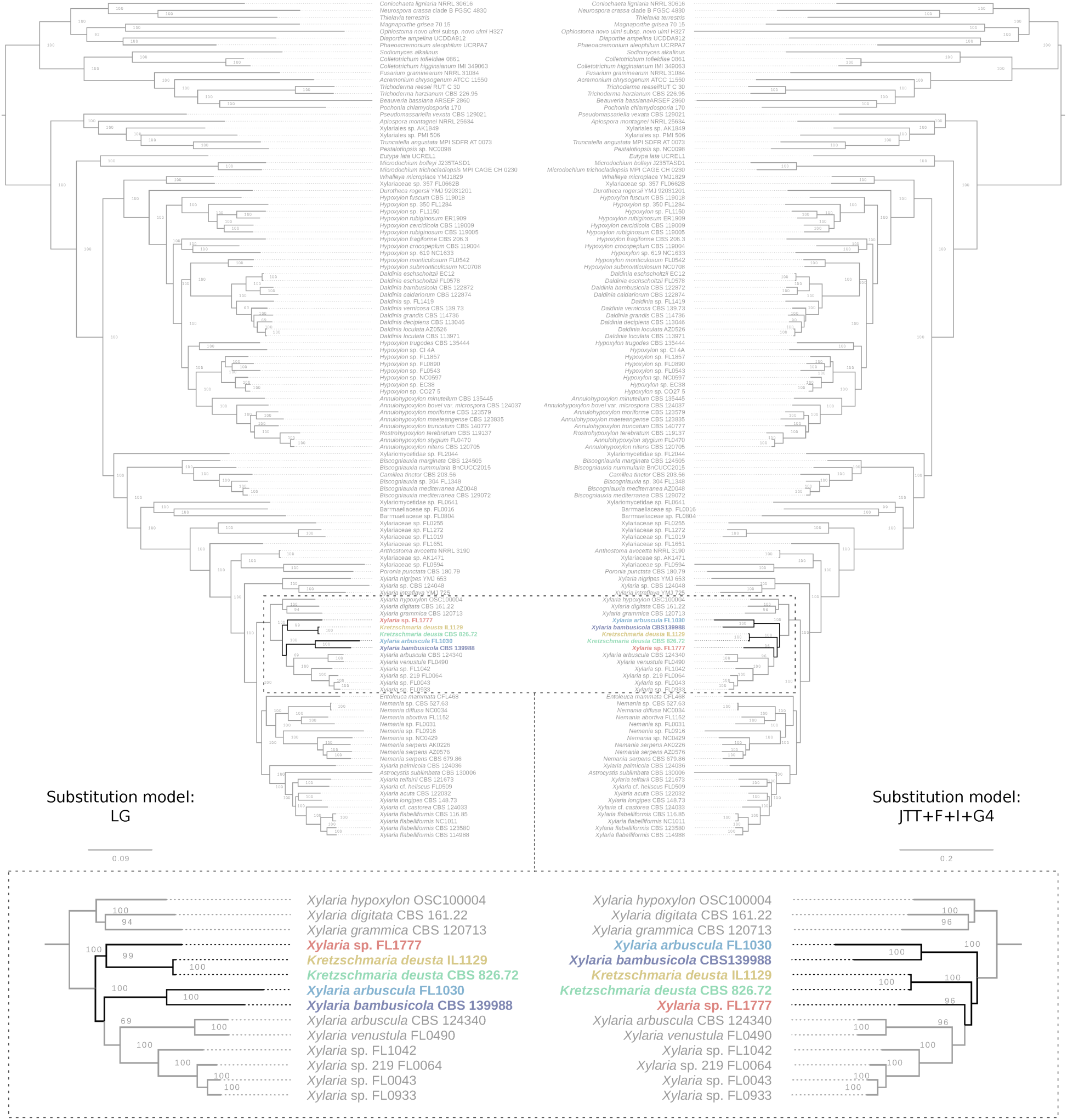
Phylogenetic tree topology was robust to outgroup taxon selection, gene set, or model of evolution. (**a**) Phylogenetic tree from the concatenated analysis of 1,526 single-copy orthogroups performed in IQ-TREE with the LG model of evolution (i.e., analysis 1; see also Supplementary Fig. 1); (**b**) Phylogenetic tree resulting from analysis of the same orthogroups, but with the JTT + F + I + G4 model of evolution (i.e., analysis 2). Topological conflicts were rare; however, the placement of five Xylariaceae s.l. taxa differed slightly with different models of evolution.

**Supplementary Figure 13.**
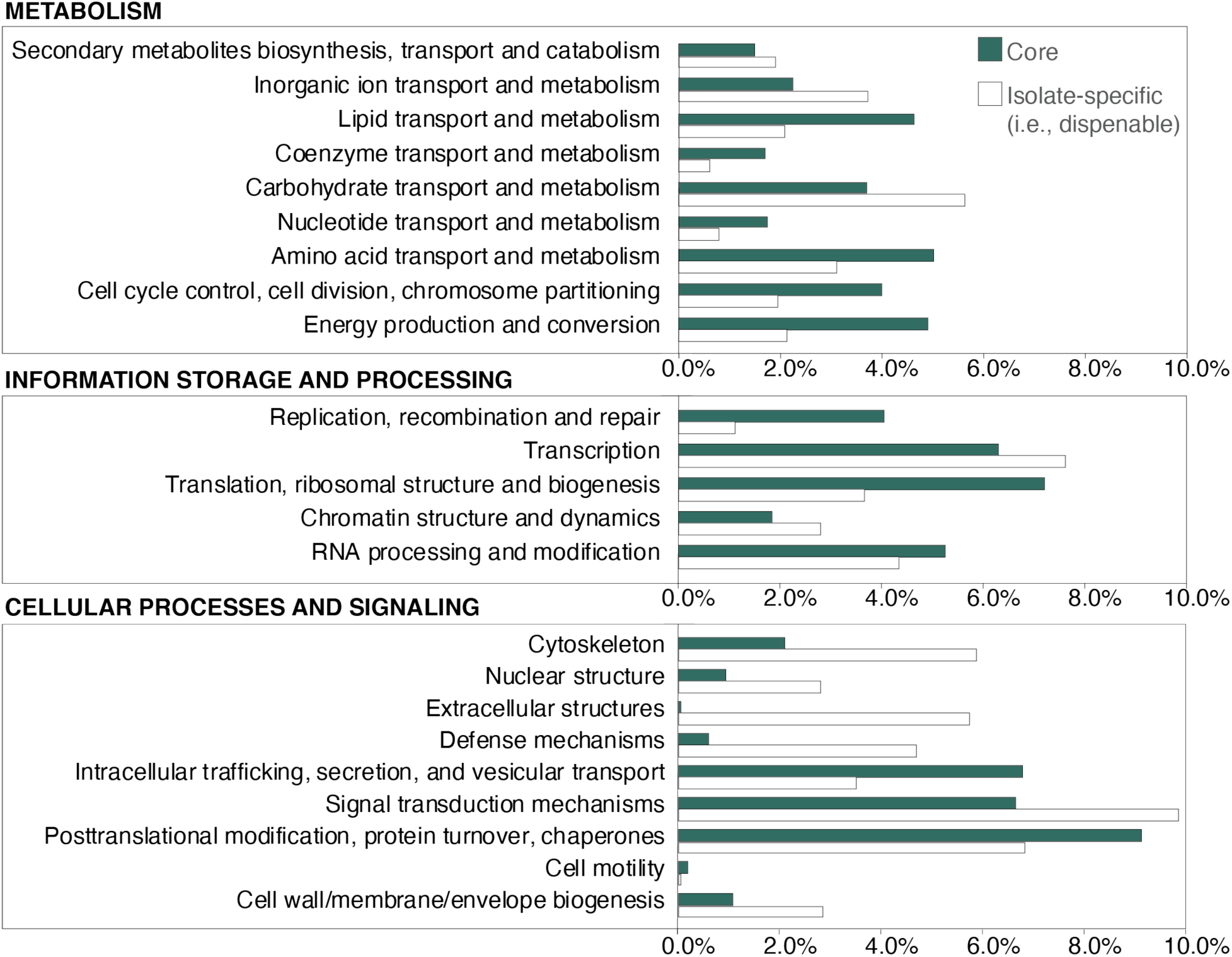
Comparison of functional annotations for core and dispensable orthogroups. Bar graphs showing the relative abundance of different functional categories represented by “core” vs. “dispensable” orthogroups. Orthogroups were annotated with euKaryotic Orthologous Groups (KOGs; see Supplementary Table 7f).

**Supplementary Figure 14.**
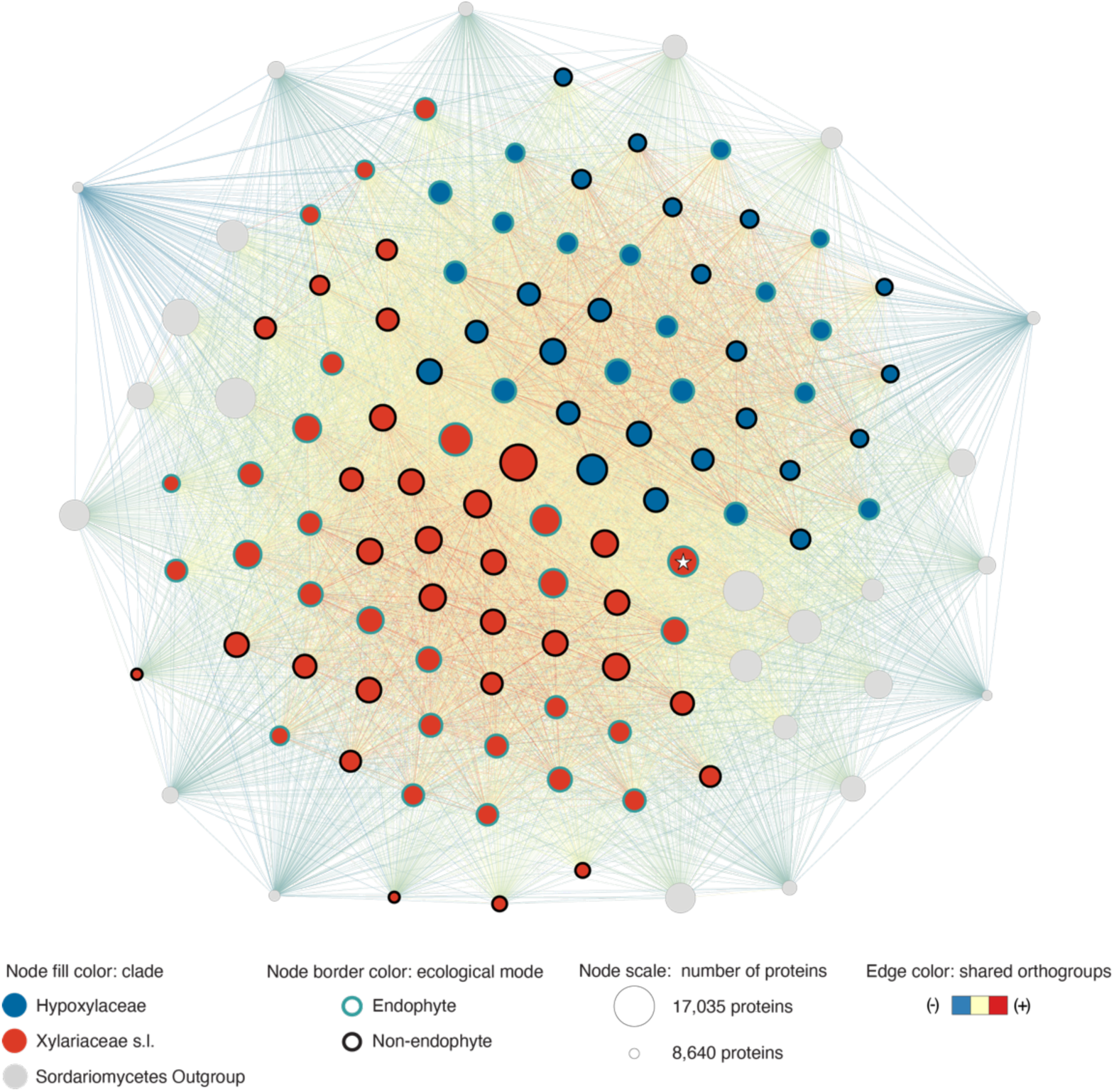
Network analysis of individual proteomes illustrates the importance of major clade affiliation. Proteomes are represented by nodes, scaled by the count of proteins, colored by clade (fill) and ecological mode (border), and positioned by a force-directed layout algorithm. Edges between two nodes are weighted by the number of shared orthogroups. The node with a star represents Xylariaceae sp. FL2044.

**Supplementary Figure 15.**
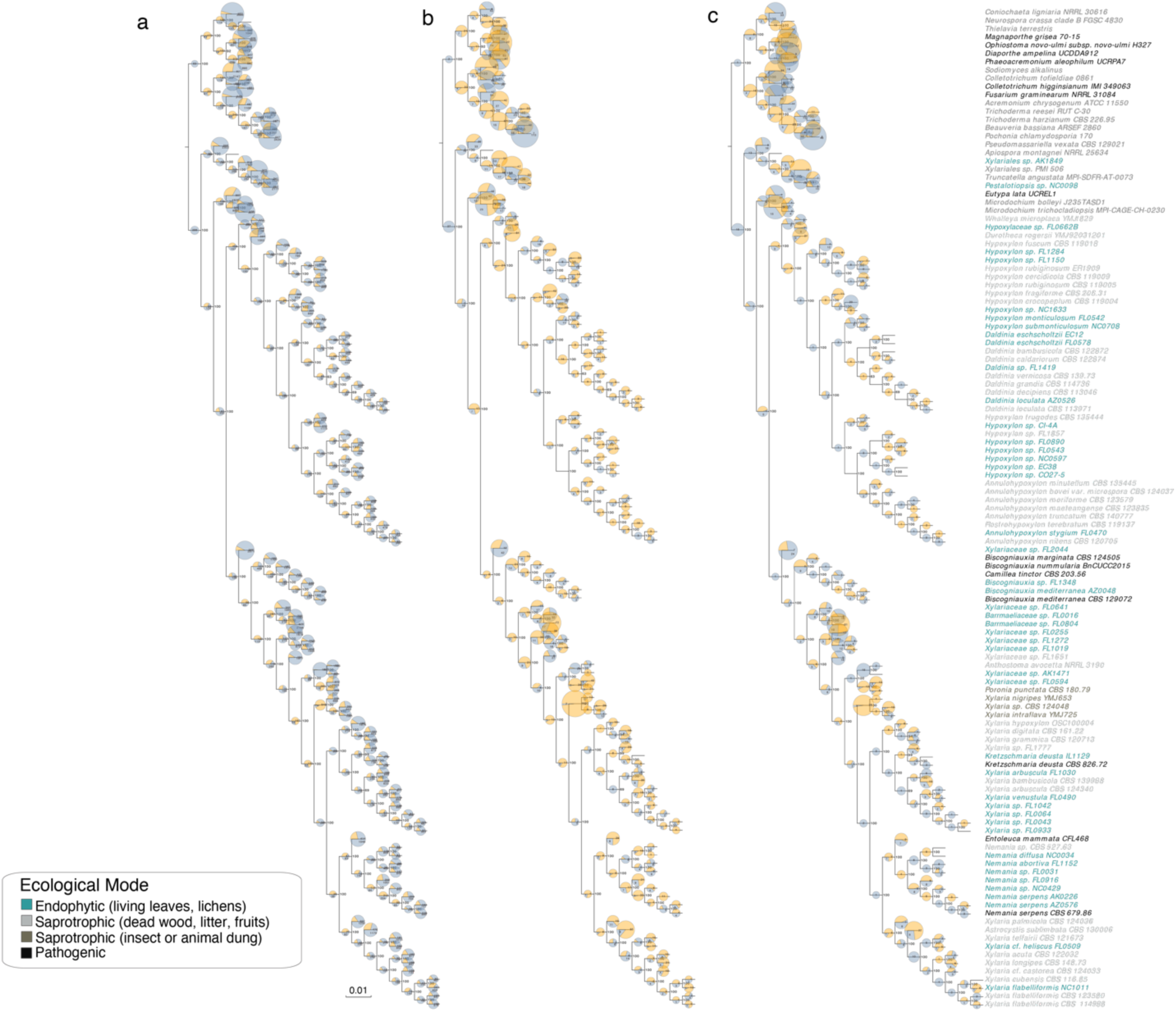
Ancestral state reconstruction of orthogroups. The number of orthogroup gain (blue) and loss (orange) events for each node (inferred using the asymmetric Wagner parsimony method: gap penalty = 1) are shown on the ML phylogenomic tree. The size of each pie chart is proportional to the total number of events inferred along the branch. Reconstructions were performed for (**a**) all orthogroups; (**b**) orthogroups annotated as CAZymes; and (**c**) orthogroups annotated as PCWDEs. Taxon names are colored by ecological mode (see legend). Predicted gains and losses were visualized on the phylogeny using EvolView^133^.

**Supplementary Figure 16.**
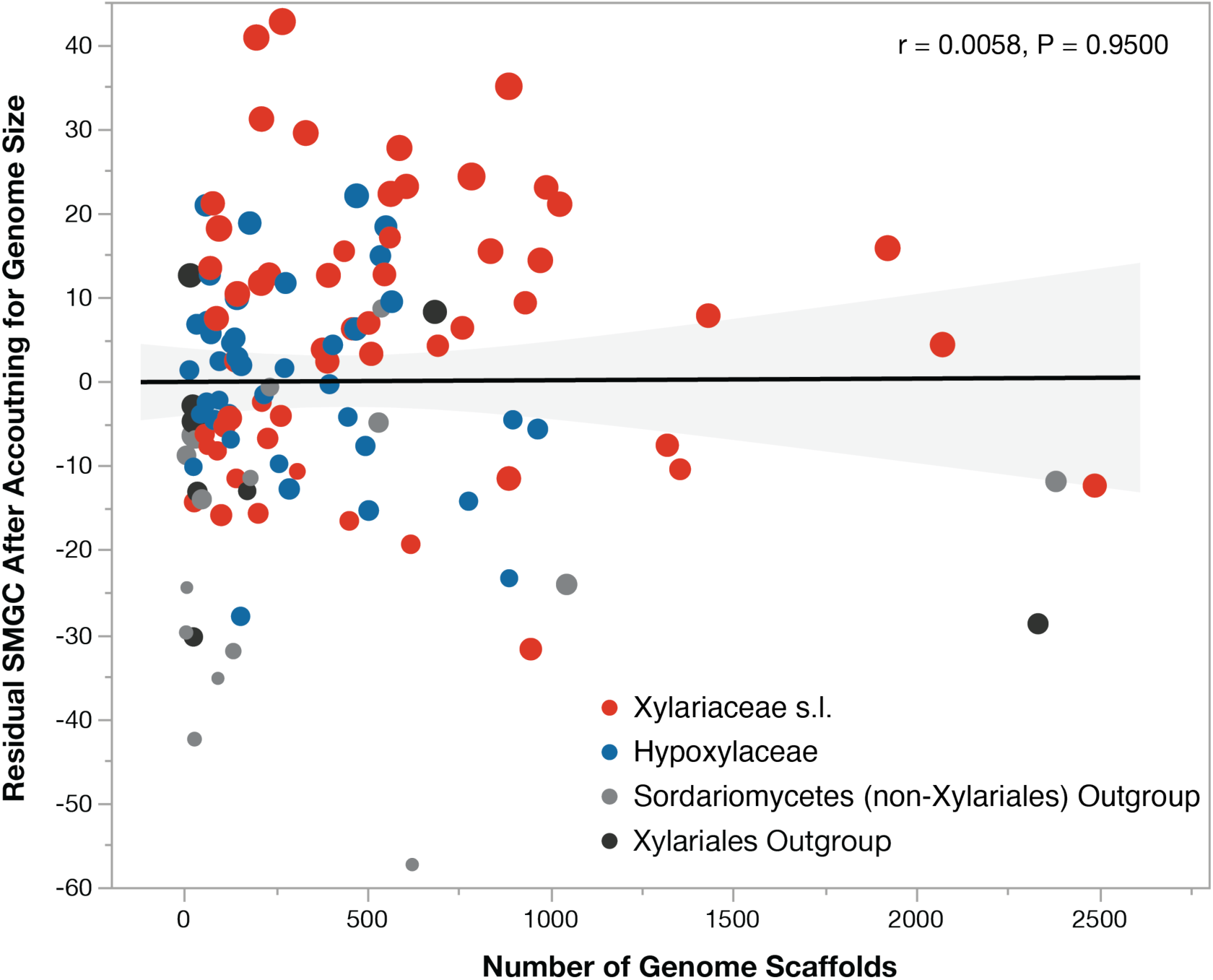
The number of predicted SMGCs is not related to genome assembly. Relationship of predicted SMGC content (residuals after accounting for genome size) and the number of scaffolds for 121 genomes. Points are colored by clade and their size is proportional to the raw number of SMGC per genome (range 16-119).

## Notes

### Competing Interest Statement

The authors have declared no competing interest.

